# Senescent cells inhibit muscle differentiation via the SASP-lipid 15d-PGJ_2_ mediated modification and control of HRas

**DOI:** 10.1101/2023.12.23.573200

**Authors:** Swarang Sachin Pundlik, Alok Barik, Ashwin Venkateshvaran, Snehasudha Subhadarshini Sahoo, Mahapatra Anshuman Jaysingh, Raviswamy G H Math, Arvind Ramanathan

## Abstract

Senescent cells, which are characterized by multiple features such as increased expression of Senescence-Associated β-galactosidase activity (SA β-gal) and cell cycle inhibitors such as p21 or p16, accumulate with tissue damage and dysregulate tissue homeostasis. In the context of skeletal muscle, it is known that agents used for chemotherapy such as Doxorubicin cause buildup of senescent cells, leading to the inhibition of tissue regeneration. Senescent cells influence the neighboring cells via numerous secreted factors which form the senescence-associated secreted phenotype (SASP). Lipids are emerging as a key component of SASP that can control tissue homeostasis. Arachidonic acid-derived lipids have been shown to accumulate within senescent cells, specifically 15d-PGJ_2_, which is an electrophilic lipid produced by the non-enzymatic dehydration of the prostaglandin PGD_2_. In this study, we show that 15d-PGJ_2_ is also released by Doxorubicin-induced senescent cells as a SASP factor. Treatment of skeletal muscle myoblasts with the conditioned medium from these senescent cells inhibits myoblast fusion during differentiation. Inhibition of L-PTGDS, the enzyme that synthesizes PGD_2_, diminishes the release of 15d-PGJ_2_ by senescent cells and restores muscle differentiation. We further show that this lipid post-translationally modifies Cys184 of HRas in skeletal muscle cells, causing a reduction in the localization of HRas to the Golgi, increased HRas binding to RAF RBD, and activation of cellular MAPK-Erk signaling (but not the Akt signaling). Mutating C184 of HRas prevents the ability of 15d- PGJ_2_ to inhibit the differentiation of muscle cells and control the activity of HRas. This work shows that 15d-PGJ_2_ released from senescent cells could be targeted to restore muscle homeostasis after chemotherapy.

## Introduction

Senescent cells are important drivers of aging and damage-associated loss of tissue homeostasis(Childs et al., 2015). Anti-cancer chemotherapy presents an important context where treatment with chemotherapeutics such as Doxorubicin (Doxo) causes widespread cellular senescence which inhibits tissue homeostasis and regeneration, including in skeletal muscles(Francis et al., 2022). It has been shown that Doxo causes systemic inflammation and leads to the emergence of senescent cells across tissues(Di Leonardo et al., 1994; Hu and Zhang, 2019; Robles and Adami, 1998). Senescent cells negatively affect tissue homeostasis and regeneration by releasing factors including proteins like growth factors, matrix metalloproteases, cytokines, and chemokines, and small molecules like fatty acid derivatives(Campisi, 2005; Coppé et al., 2010; Dilley et al., 2003; Krtolica et al., 2001; Parrinello et al., 2005; Shelton et al., 1999; Yang et al., 2006). The release of these factors from senescent cells is called the Senescence-Associated Secretory Phenotype (SASP). It is expected that these SASP factors and their mechanisms of action will vary depending on cellular and tissue contexts. Identifying SASP factors and their underlying mechanistic targets will be critical for building an understanding of how senescent cells control tissue homeostasis(Coppé et al., 2010; Davalos et al., 2010). Lipids are a less explored family of SASP factors, and it is important to understand how they affect tissue regeneration(Hamsanathan and Gurkar, 2022). We have previously shown that senescent cells have increased intracellular levels of prostaglandin 15d-PGJ_2_(Wiley et al., 2021), a non-enzymatic dehydration product of prostaglandin PGD_2_(Shibata et al., 2002). In the context of skeletal muscle, PGD_2_ and 15d-PGJ_2_ have been shown to negatively regulate muscle differentiation via mechanisms that do not depend on a cognate receptor(Hunter et al., 2001; Veliça et al., 2010). Here we study the role of 15d-PGJ_2_ as a member of the SASP and identify the mechanisms by which it might negatively affect muscle regeneration. 15d-PGJ_2_ has been previously shown to covalently modify multiple proteins like MAPK1, MCM4, EIF4A-I, PKM1, GFAP etc. in endothelial and neuronal cells(Marcone and Fitzgerald, 2013; Yamamoto et al., 2011). 15d-PGJ_2_ was shown to be covalently modifying HRas in NIH3T3, Cos1, and IMR90 cell lines(Luis Oliva et al., 2003; Wiley et al., 2021). We further studied HRas as an important target that might mediate the effects of 15d-PGJ_2_ on muscle differentiation via covalent modification. We investigated HRas as a possible effector of 15d-PGJ_2_ because (i) HRas belongs to the Ras superfamily of small molecule GTPases and is a known regulator of key cellular processes(Davis et al., 1983; Harvey, 1964; Kirsten and Mayer, 1967; Vetter and Wittinghofer, 2001). (ii) constitutively active HRas mutant (HRas V12) has been shown to inhibit the differentiation of myoblasts by inhibiting MyoD and Myogenin expression(Konieczny et al., 1989; Lassar et al., 1989; Olson,’ et al., 1987; Van Der Burgt et al., 2007). (iii) Downstream signaling of HRas is important for muscle homeostasis as skeletal and cardiac myopathies are observed in individuals carrying constitutively active mutants of HRas(Engler et al., 2021; Konieczny et al., 1989; Lee et al., 2010; Olson,’ et al., 1987; Scholz et al., 2009; Van Der Burgt et al., 2007). HRas is highly regulated by lipid modifications, it undergoes reversible palmitoylation and de-palmitoylation at C-terminal cysteines, which regulate the intracellular distribution and activity of HRas(Gutierrez et al., 1989; Lu and Hofmann, 1995; Rocks et al., 2005). In this study, we show that 15d-PGJ_2_ is synthesized and released by senescent myoblasts upon treatment with Doxo. 15d-PGJ_2_, taken up by the myoblasts, covalently modifies HRas at cysteine 184 and activates it. We also show that previously reported inhibition of differentiation of myoblasts by 15d-PGJ_2_ depends on HRas C-terminal cysteines, notably cysteine 184. This study provides a mechanism by which prostaglandins secreted as SASP inhibit the differentiation of myoblasts, affecting muscle homeostasis in patients undergoing chemotherapy.

## Results

### Doxorubicin (Doxo) treatment induces senescence in mouse skeletal muscles and C2C12 mouse myoblasts

Doxorubicin-mediated DNA damage has been shown to induce senescence in cells(Di Leonardo et al., 1994; Hu and Zhang, 2019; Robles and Adami, 1998). Therefore, we injected B6J mice intraperitoneally with Doxorubicin (Doxo) (5mg/kg) every 3 days for 9 days and observed induction of DNA damage-mediated senescence in hindlimb skeletal muscles (Fig. S1A). We observed an increase in the expression of p21 and increased nuclear levels of the DNA damage marker γH2A.X in mouse Gastrocnemius muscles (Fig. 1A and B). We also observed a significant increase in the mRNA levels of known senescence markers (p16 and p21), SASP factors (CXCL1, CXCL2, TNF1α, IL6, TGFβ1) in skeletal muscles of mice treated with Doxo compared to that of mice treated with saline (Fig. 1C). These observations suggest that there is induction of senescence in skeletal muscles of mice upon treatment with Doxo.

**Figure 1.**
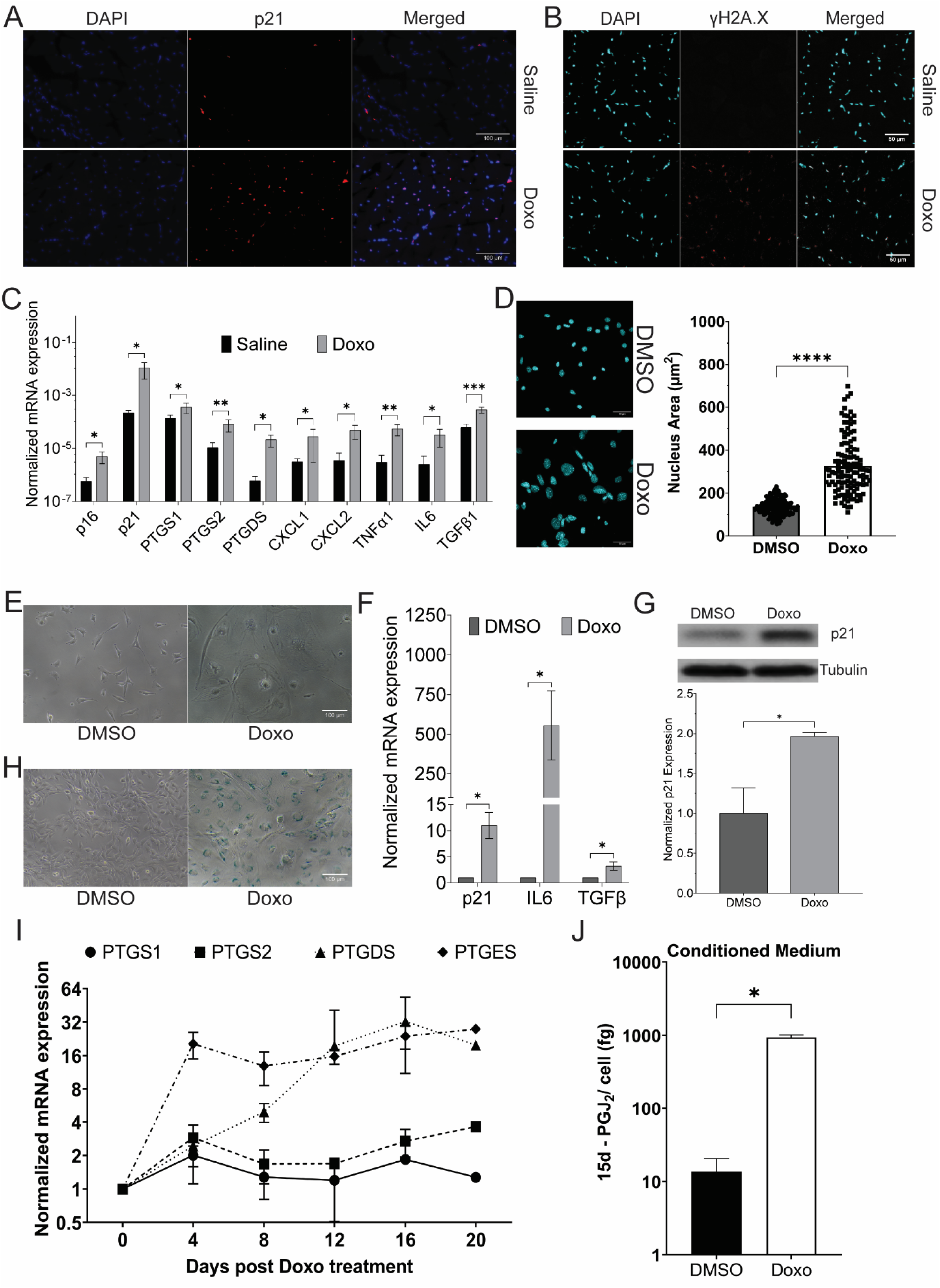
Prostaglandin synthesis and release by Doxo-induced senescent cells inhibits myoblast differentiation. A. Expression and localization of tumor suppressor protein p21, measured by immunofluorescence, in hindlimb skeletal muscles of mice after 11 days of treatment with Doxo (5 mg/kg) or Saline. B. Representative confocal micrograph of expression of γH2A.X in the gastrocnemius muscle of mice treated with Doxo (5 mg/kg) or Saline. C. Expression of mRNAs of senescence markers (p16 and p21), SASP factors (CXCL1, CXCL2, TNFα1, IL6, TGFβ1), and enzymes involved in the biosynthesis of prostaglandin PGD_2_/15d-PGJ_2_ (PTGS1, PTGS2, PTGDS), measured by qPCR, in hindlimb skeletal muscles of mice after 11 days of treatment with Doxo (5 mg/kg) or Saline. D. A representative confocal micrograph and a scatter plot of the nuclear area of C2C12 myoblasts, measured by immunofluorescence, after 16 days of treatment with Doxo (150 nM) or DMSO. E. A representative widefield micrograph of cell morphology in C2C12 myoblasts after 16 days of treatment with Doxo (150 nM) or DMSO. F. Expression of mRNA of cell cycle inhibitor p21 and SASP factors (IL6 and TGFβ), measured by qPCR, in C2C12 myoblasts after 16 days of treatment with Doxo (150 nM) or DMSO. G. Expression of cell cycle inhibitor p21, measured by immunoblot, in C2C12 myoblasts after 16 days of treatment with Doxo (150 nM) or DMSO. H. Activity of Senescence Associated β-galactosidase (SA β-gal), measured by X-gal staining at pH∼6, in C2C12 myoblasts after 16 days of treatment with Doxo (150 nM) or DMSO. I. Expression of mRNAs of prostaglandin biosynthetic enzymes, measured by qPCR, in C2C12 myoblasts after treatment with Doxo (150 nM) or DMSO. J. Concentration of 15d-PGJ_2_ released from quiescent or senescent C2C12 cells. (Statistical significance was tested by the two-tailed student’s t-test ns=p>0.05, *=p<0.05, **=p<0.01, ***=p<0.001, ****=p<0.0001)

C2C12 cells have been shown to undergo senescence after DNA damage, as assessed by an increase in the levels of SA β-gal and known markers of SASP (IL1α, IL6, CCL2, CXCL2, CXCL10)(Moiseeva et al., 2022). We treated C2C12 myoblasts with Doxo (150 nM) and observed a significant increase in the size of the nuclei (Fig. 1D), flattened cell morphology with an increase in the cell size (Fig. 1E), a significant increase in the mRNA levels of cell cycle inhibitor p21 and SASP factors IL6 and TGFβ (Fig. 1F), a significant increase in the protein levels of p21 (Fig. 1G), and an increase in the levels of SA β-gal (Fig. 1H) in C2C12 cells treated with Doxo. These observations suggest that C2C12 cells undergo senescence upon treatment with Doxo.

### Doxo-mediated senescence induces synthesis and release of 15d-PGJ_2_ in C2C12 myoblasts and mouse skeletal muscle

Synthesis of prostaglandins by senescent cells has previously been reported(Wiley et al., 2021; Wiley and Campisi, 2021). Specifically, levels of PGD_2_ and its metabolite 15d-PGJ_2_ have been shown to be significantly increased in senescent cells. Therefore, we measured the levels of mRNA of enzymes involved in the synthesis of PGD_2_/15d-PGJ_2_ (PTGS1, PTGS2, and PTGDS), in the gastrocnemius muscle of mice after treatment with Doxo. We observed a significant increase in the mRNA levels of PTGS1, PTGS2, and PTGDS enzymes in the skeletal muscle of mice treated with Doxo (Fig. 1C). We also observed a time-dependent increase in the mRNA levels of PTGS1, PTGS2, PTGDS, and PTGES enzymes in C2C12 cells treated with Doxo compared to Day 0 (Fig. 1I). Expression of enzyme PTGES was elevated on Day 4, whereas the expression of Prostaglandin D synthase (PTGDS) increased only after Day 8, reaching maximum expression on Day 12. These observations suggest an increase in the synthesis of prostaglandins in senescent cells.

15d-PGJ_2_ is a non-enzymatic dehydration product of PGD_2_(Shibata et al., 2002). We observed an increase in the mRNA levels of synthetic enzymes of 15d-PGJ_2_ in senescent C2C12 cells. Therefore, we measured the levels of 15d-PGJ_2_ released by senescent C2C12 cells using targeted mass spectrometry (Fig. S1C, D, E, and F). The concentration of 15d-PGJ_2_ was quantified by monitoring the transition of the m/z of ions from 315.100 → 271.100 using a SCIEX 6500 mass spectrometer. We plotted a standard curve using purified 15d-PGJ_2_ (Fig. S1F) to quantify the concentration of 15d-PGJ_2_. We used the representative peaks from the conditioned medium collected from C2C12 cells incubated in 0.2% serum medium for 3 days (Quiescent cells) and C2C12 cells treated with Doxo (150 nM) (Senescent cells) to measure the concentration of 15d-PGJ_2_ released by quiescent cells or senescent C2C12 cells. We observed a significant increase (∼100 fold) in the concentration of 15d-PGJ_2_ in the conditioned medium from senescent cells as compared to that in quiescent cells (Fig. 1J). This suggests that senescent C2C12 cells release 15d-PGJ_2_ in the medium.

### Prostaglandin PGD_2_ and its metabolites in the conditioned medium of senescent cells inhibit the differentiation of C2C12 myoblasts

15d-PGJ_2_ (the final non-enzymatic dehydration product of PGD_2_) has been shown to inhibit the differentiation of myoblasts(Hunter et al., 2001). We observed the release of 15d-PGJ_2_ by senescent cells, showing that 15d-PGJ_2_ is a SASP factor (Fig. 1F). Conditioned medium of senescent cells inhibits the differentiation of myoblasts in myotonic dystrophy type 1(Conte et al., 2023). Therefore, we tested whether 15d-PGJ_2_, the terminal dehydration product of PGD_2_, is required for the inhibitory effect of SASP on the differentiation of myoblasts. We treated C2C12 myoblasts with the conditioned medium of senescent cells or senescent cells treated with 30 µM of AT-56 (a well-characterized inhibitor of prostaglandin D synthase (PTGDS))(Hu et al., 2021; S. Hu et al., 2023; Shunfeng Hu et al., 2023; Irikura et al., 2009) and measured the differentiation of myoblasts by calculating the fusion index. We observed a significant decrease (∼20%) in the fusion index of the C2C12 myoblasts treated with the conditioned medium of senescent cells (Fig. 2A), suggesting that SASP factors decrease the differentiation of myoblasts. This decrease in the inhibition was rescued in myoblasts treated with the conditioned medium of senescent cells treated with AT-56 (Fig. 2A). This suggests that prostaglandins PGD_2_/15d-PGJ_2_ released by senescent cells as SASP factors can inhibit the differentiation of myoblasts.

**Figure 2.**
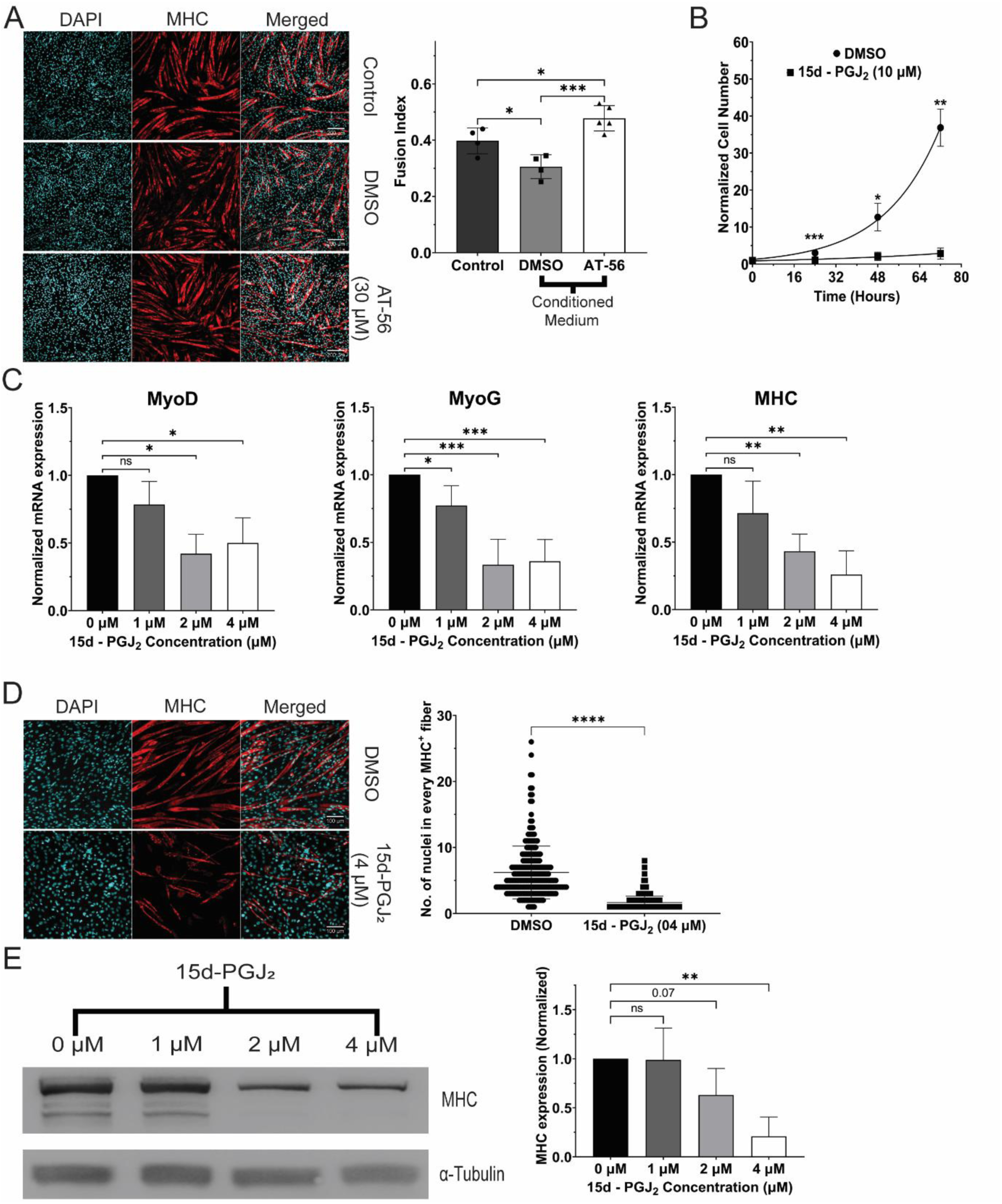
15d-PGJ_2_ inhibits differentiation of myoblasts. A. Expression of MHC protein and the fusion of myoblasts in myotubes, measured by immunofluorescence, after treatment with conditioned medium of senescent cells treated with PTGDS inhibitor AT-56 (30 µM) or DMSO B. Normalized number of C2C12 myoblasts treated with 15d-PGJ_2_ (10 µM) or DMSO C. Expression of mRNAs of markers of differentiation (MyoD, MyoG, and MHC), measured by qPCR, in C2C12 myoblasts treated with 15d-PGJ_2_ (1 µM, 2 µM, or 4 µM) or DMSO. D. Expression of MHC protein and the fusion of myoblasts in syncytial myotubes, measured by immunofluorescence, after treatment with 15d-PGJ_2_ (4 µM) or DMSO. E. Expression of MHC protein, measured by immunoblotting, in primary human skeletal myoblasts after treatment with 15d-PGJ_2_ (1 µM, 2 µM, or 4 µM) or DMSO for 5 days. (Statistical significance was tested by the two-tailed student’s t-test ns=p>0.05, *=p<0.05, **=p<0.01, ***=p<0.001, ****=p<0.0001)

### 15d-PGJ2 inhibits the proliferation and differentiation of mouse and human myoblasts

15d-PGJ_2_ has been shown to affect the proliferation of cancer cell lines, both positively and negatively (Chen et al., 2003; Choi et al., 2020; Slanovc et al., 2024; Yen et al., 2014). We measured the effect of 15d-PGJ_2_ on the proliferation of C2C12 myoblasts. We treated C2C12 myoblasts with 15d-PGJ_2_ (10 µM) or DMSO in DMEM 10% Serum medium for 72 hours and observed a significant decrease in the proliferation of C2C12 cells after treatment with 15d-PGJ_2_ (Fig. 2B). The doubling time of C2C12 cells was also increased upon treatment with 15d-PGJ_2_ (57.24 hours) compared to DMSO (13.76 hours). This suggests that 15d-PGJ_2_ decreases the proliferation of C2C12 myoblasts.

We measured the differentiation of C2C12 mouse and primary human myoblasts after treatment with 15d-PGJ_2_. To rule out the toxic effects of 15d-PGJ_2_ on cell physiology, we treated C2C12 cells with 15d-PGJ_2_ (1 µM, 2 µM, 4 µM, 5 µM, and 10 µM) in the C2C12 differentiation medium, and measured the viability of cells after 24 hours of treatment, using an MTT viability assay. We judged that 15d-PGJ_2_ was not cytotoxic up to 5 µM in the C2C12 differentiation medium (Fig. S2A). Based on this, we treated differentiating myoblasts with 15d-PGJ_2_ (1 µM, 2 µM, and 4 µM) for 5 days to measure the effects of 15d-PGJ_2_ treatment on differentiation of myoblasts. We observed a dose-dependent decrease in the mRNA levels of MyoD, MyoG, and MHC in differentiating C2C12 cells after treatment with 15dPGJ_2_ (Fig. 2C). There was a significant decrease in the no. of nuclei in individual MHC^+ve^ fiber (∼75%) in C2C12 cells treated with 15d-PGJ_2_ (4 µM) compared to DMSO (Fig. 2D), suggesting a decrease in the fusion of myoblasts in myotubes. We also observed a dose-dependent decrease in the protein levels of MHC in differentiating primary human myoblasts upon treatment with 15d-PGJ_2_ (Fig. 2E). Together, these observations suggest that 15d-PGJ_2_ inhibits the differentiation of both mouse and human myoblasts.

### Biotinylated 15d-PGJ_2_ covalently modifies HRas at Cysteine 184

15d-PGJ_2_ has been shown to covalently modify several proteins including p53 and NF-κB, which are involved in several key biological processes(Marcone and Fitzgerald, 2013). HRas was identified to be covalently modified by 15d-PGJ_2_ at cysteine 184 in NIH3T3 and Cos1 cells(Luis Oliva et al., 2003). Therefore, we tested whether 15d-PGJ_2_ could covalently modify HRas in C2C12 cells. We treated C2C12 cells expressing the EGFP-tagged wild-type HRas with biotinylated 15d-PGJ_2_ (5 µM). We then immunoprecipitated biotinylated 15d-PGJ_2_ using streptavidin. We observed a significant increase (∼3.5 fold) in the pulldown of HRas upon treatment with 15d-PGJ_2_biotin compared to DMSO (Fig. 3A), suggesting an interaction between 15d-PGJ_2_ and HRas. To measure the role of individual C-terminal cysteines in the binding of HRas with 15d-PGJ_2_, we treated C2C12 cells expressing the EGFP-tagged C181S and C184S mutants of HRas with biotinylated 15d-PGJ_2_ (5 µM), and immunoprecipitated using streptavidin. We observed that the intensity of EGFP-tagged HRas was significantly decreased in cells expressing the C184S mutant (∼80% decrease compared to HRas WT) but not in those expressing the C181S mutant (Fig. 3A). This suggests that 15d-PGJ_2_ covalently modifies HRas at cysteine 184 in C2C12 cells.

**Figure 3.**
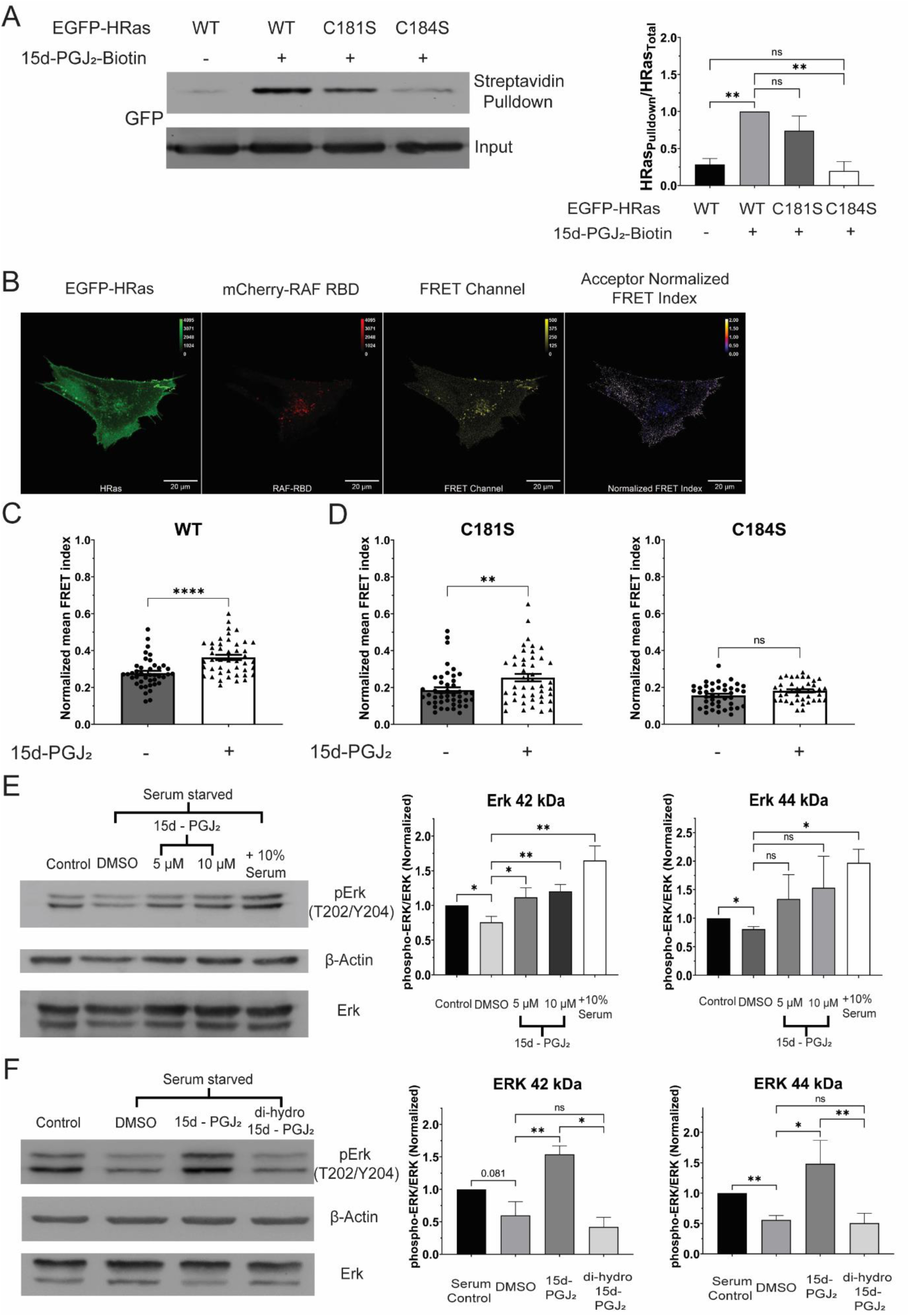
15d-PGJ_2_ covalently modifies HRas at Cysteine 184 and activates the HRas-MAPK pathway via the electrophilic cyclopentenone ring. A. Streptavidin-immunoprecipitation of EGFP-HRas, measured by immunoblotting, in C2C12 cells after 3 hours of treatment with 15d-PGJ_2_-Biotin (5 µM). B. Representative confocal micrograph of Fluorescence Resonance Energy Transfer (FRET) between EGFP-tagged HRas (EGFP-HRas) and mCherry-tagged Ras binding domain (RBD) of RAF kinase (mCherry-RAF RBD). C. Activation of the EGFP-tagged wild type HRas (HRas WT), measured by FRET, before and after 1 hour of treatment with 15d-PGJ_2_ (10 µM) after starvation for 24 hours. D. Activation of the EGFP-tagged C-terminal cysteine mutants of HRas (HRas C181S and HRas C184S), measured by FRET, before and after 1 hour of treatment with 15d-PGJ_2_ (10 µM) after starvation for 24 hours. E. Phosphorylation of Erk (42 kDa and 44 kDa), measured by immunoblotting, in C2C12 cells after 1 hour of treatment with 15d-PGJ_2_ (5 µM, 10 µM) or DMSO after starvation for 24 hours. F. Phosphorylation of Erk (42 kDa and 44 kDa), measured by immunoblotting, in C2C12 cells after 1 hour of treatment with 15d-PGJ_2_ (10 µM)/ 9,10-dihydro-15d-PGJ_2_ (10 µM) or DMSO after starvation for 24 hours. (Statistical significance was tested by the two-tailed student’s t-test ns=p>0.05, *=p<0.05, **=p<0.01, ***=p<0.001, ****=p<0.0001)

### 15d-PGJ2 increases the FRET between EGFP-HRas and mCherry-RAF-RBD in wild-type and C181S mutant but not in the C184S mutant of HRas

We next tested the effect of covalent modification of HRas by 15d-PGJ_2_ on HRas GTPase activity using FRET. mCherry-RAF-RBD is a well-characterized sensor of the activity of HRas. RAF-RBD binds to the activated HRas upon activation of HRas, allowing FRET between EGFP and mCherry(Rocks et al., 2005). We co-expressed EGFP-tagged HRas (EGFP-HRas) with mCherry-RAF-RBD in C2C12 myoblasts (Fig. 3B). We measured the efficiency of FRET between EGFP and mCherry using an ImageJ plugin, FRET analyzer(Hachet-Haas et al., 2006). We compared the mean acceptor normalized FRET index in C2C12 myoblasts co-expressing EGFP-HRas WT and mCherry-RAF-RBD before and after treatment of 15d-PGJ_2_ (10 µM) for 1 hour. We observed a significant increase (∼30%) in the mean acceptor normalized FRET index upon treatment with 15d-PGJ_2_ (Fig. 3C). This suggests that 15d-PGJ_2_ activates HRas. To measure the role of individual C-terminal cysteines in 15d-PGJ_2_ mediated activation of HRas, we co-expressed EGFP-HRas C181S or C184S with mCherry-RAF-RBD in C2C12 myoblasts. We measured the mean acceptor normalized FRET index before and after 1 hour of treatment with 15d-PGJ_2_ (10 µM). We observed a significant increase (∼40%) in the mean acceptor normalized FRET index in cells expressing EGFP-HRas C181S upon treatment with 15d-PGJ_2_ but not in cells expressing EGFP-HRas C184S (Fig. 3D). These observations suggest that activation of HRas by 15d-PGJ_2_ occurs in a cysteine 184 dependent manner.

### 15d-PGJ2 increases phosphorylation of Erk (Thr202/Tyr204) but not Akt (S473) in C2C12 myoblasts

HRas regulates two major downstream signaling pathways, the MAP kinase (MAPK) pathway and the PI3 kinase (PI3K) pathway(Pylayeva-Gupta et al., 2011). We tested the effects of treatment with 15d-PGJ_2_ on these two downstream signaling pathways by measuring the phosphorylation of Erk (42 kDa and 44 kDa) and Akt proteins in C2C12 cells. We treated C2C12 cells with 15d-PGJ_2_ (5 µM and 10 µM) or DMSO for 1 hr (after 24 hrs. of serum starvation) and observed a dose-dependent increase in the phosphorylation of Erk (T202/Y204) (42 kDa) but not of Erk (44 kDa) (Fig. 3E). We did not observe an increase in the phosphorylation of Akt (S473) in C2C12 cells after treatment with 15d-PGJ_2_ (Fig. S3C). These observations suggest that 15d-PGJ_2_ activates the MAPK signaling pathway, but not the PI3K signaling pathway.

15d-PGJ_2_ contains a reactive electrophilic center in its cyclopentenone ring, that can react with cysteine residues of proteins(Luis Oliva et al., 2003). We tested its role in activating the MAPK signaling pathway. We measured the phosphorylation of Erk (42kDa and 44 kDa) in C2C12 cells after treatment with cells with 9,10-dihydro-15d-PGJ_2_ (10 µM), a 15d-PGJ_2_ analog which is devoid of the electrophilic center, for 1 hr (after 24 hr. of serum starvation). We observed that the phosphorylation of Erk (42 kDa and 44 kDa) in C2C12 cells treated with 9,10-dihydro-15d-PGJ_2_ was significantly reduced (∼70%) as compared to the treatment with 15d-PGJ_2_ (Fig. 3F). This shows that 15d-PGJ_2_ activates the HRas-MAPK signaling pathway via the electrophilic center in its cyclopentenone ring.

### 15d-PGJ2 increases the localization of EGFP-tagged HRas at the plasma membrane compared to the Golgi in a C-terminal cysteine-dependent manner

15d-PGJ_2_ covalently modifies cysteine 184 and activates HRas signaling (Fig. 3). Reversible palmitoylation of cysteine 181 and cysteine 184 in the C-terminal tail of HRas regulate intracellular distribution and signaling of HRas. Inhibition of palmitoylation of the C-terminal cysteine 181, either by a palmitoylation inhibitor 2-Bromopalmitate or by mutation to serine, causes accumulation of HRas at the Golgi compared to the plasma membrane and alters activity(Rocks et al., 2005). Therefore, we tested whether the modification of 15d-PGJ_2_ alters the intracellular distribution of HRas. We co-expressed the EGFP-tagged wild type and the cysteine mutants of HRas (EGFP-HRas WT/C181S/C184S) with a previously reported marker of Golgi(Shaner et al., 2008) in C2C12 cells and stained the cells with plasma membrane marker WGA-633 (Fig. 4A). We compared R_mean_, the ratio of mean EGFP-HRas intensity at the Golgi to the mean HRas intensity at the plasma membrane, to measure the distribution of HRas between the plasma membrane and the Golgi. We measured the intracellular distribution of HRas between the Golgi and the plasma membrane in C2C12 cells after treatment with 15d-PGJ_2_ (10 µM) for 24 hours in DMEM 10% serum medium and observed a significant decrease (∼20%) in the R_mean_ of C2C12 cells expressing the wild-type HRas after treatment with 15d-PGJ_2_ (Fig. 4B). However, we did not observe a change in the R_mean_ of C2C12 cells expressing HRas C181S or HRas C184S after treatment with 15d-PGJ_2_ (Fig. 4C). These observations suggest that 15d-PGJ_2_ increases the localization of HRas at the plasma membrane as compared to that in the Golgi in an HRas C-terminal cysteine-dependent manner.

**Figure 4.**
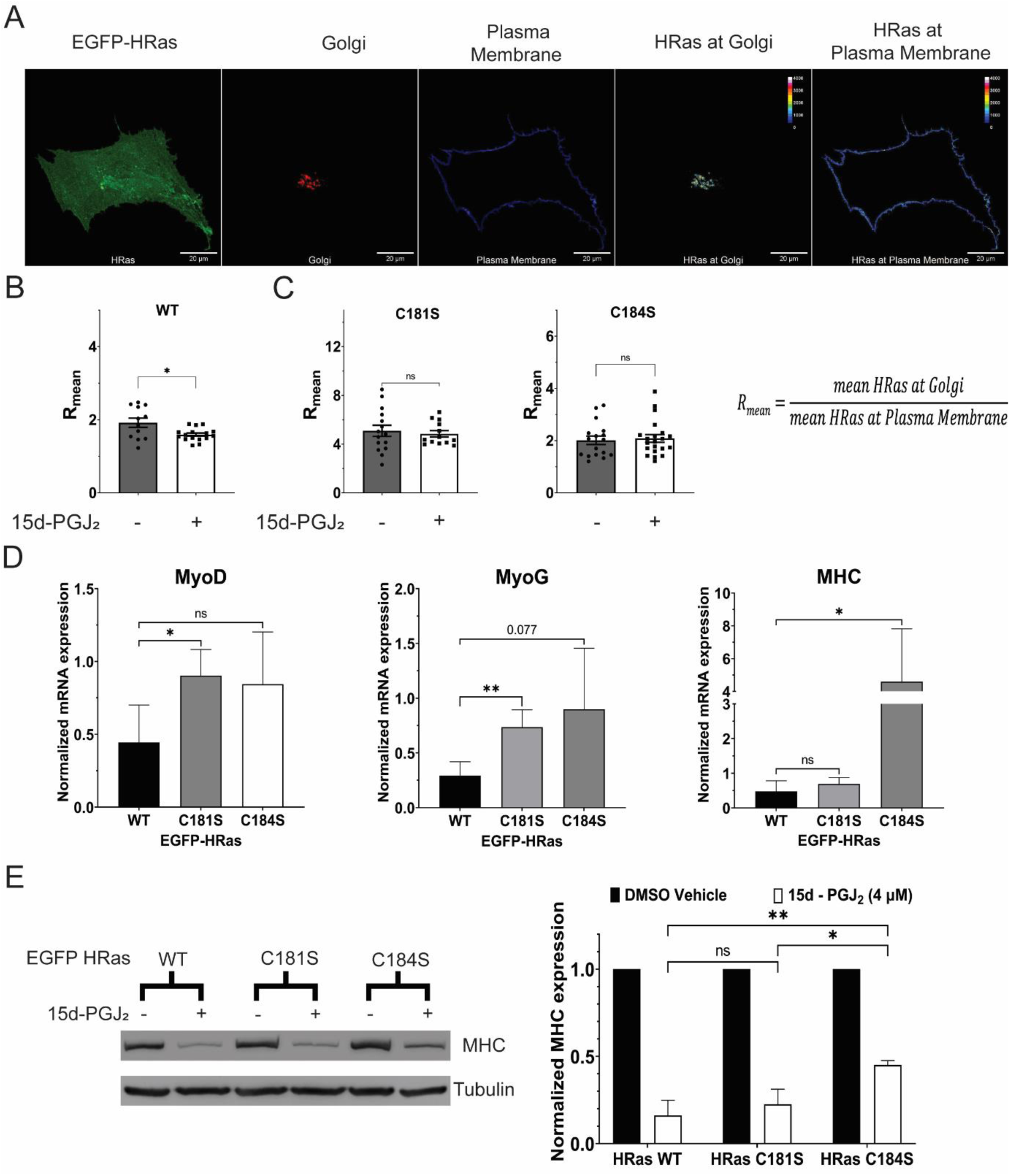
15d-PGJ_2_ controls the intracellular distribution of HRas and differentiation of C2C12 cells in an HRas C-terminal cysteine-dependent manner. A. Representative confocal micrograph of C2C12 myoblasts showing localization of EGFP-tagged HRas between the plasma membrane (stained with Alexa Fluor 633 conjugated Wheat Germ Agglutinin) and the Golgi (labelled with TagRFP-tagged Golgi resident GalT protein). A statistic R_mean_ was defined as the ratio of mean HRas intensity at the Golgi to the mean HRas intensity at the plasma membrane. B. Distribution of the wild-type HRas between the Golgi and the plasma membrane, measured by R_mean_, in C2C12 myoblasts treated with 15d-PGJ_2_ (10 µM) or DMSO for 24 hours. C. Distribution of the C-terminal cysteine mutants of HRas between the Golgi and the plasma membrane, measured by R_mean_, in C2C12 myoblasts treated with 15d-PGJ_2_ (10 µM) or DMSO for 24 hours. D. Expression of mRNAs of known markers of differentiation (MyoD, MyoG, and MHC), measured by qPCR, in differentiating C2C12 myoblasts expressing the EGFP-tagged wild-type and the C-terminal cysteine mutants of HRas after treatment with 15d-PGJ_2_ (4 µM) or DMSO for 5 days. E. Expression of MHC protein, measured by immunoblotting, in differentiating C2C12 myoblasts expressing the EGFP-tagged wild-type and the C-terminal cysteine mutants of HRas after treatment with 15d-PGJ_2_ (4 µM) or DMSO for 5 days. (Statistical significance was tested by the two-tailed student’s t-test ns=p>0.05, *=p<0.05, **=p<0.01, ***=p<0.001, ****=p<0.0001)

### 15d-PGJ2 mediated inhibition of differentiation of C2C12 cells is rescued by C181S and C184S mutants of HRas

HRas inhibits the differentiation of C2C12 myoblasts(Engler et al., 2021; Konieczny et al., 1989; Lassar et al., 1989; Lee et al., 2010; Olson,’ et al., 1987; Scholz et al., 2009; Van Der Burgt et al., 2007).15d-PGJ_2_ covalently modifies cysteine 184 and activates HRas (Fig. 3). Therefore, we tested whether the inhibition of myoblast differentiation by 15d-PGJ_2_ depends on the activation of HRas signaling by modification of the C-terminal cysteine 184. We expressed the wild-type and the cysteine mutants of HRas (EGFP-HRas WT/C181S/C184S) in C2C12 myoblasts and treated the cells with 15d-PGJ_2_ (4 µM) or DMSO during differentiation. We observed a decrease in the levels of mRNA of MHC in C2C12 cells expressing HRas WT and HRas C181S after 5 days of treatment with 15d-PGJ_2_. We did not observe this in expressing HRas C184S (Fig. 4D). We also observed a significant decrease in the protein levels of MHC in differentiating C2C12 cells expressing HRas WT and HRas C181S after treatment with 15d-PGJ_2_ (Fig. 4E). This decrease was partially rescued in cells expressing HRas C184S (Fig. 4E). These observations suggest that the inhibition of myoblast differentiation by 15d-PGJ_2_ depends on modification of HRas C-terminal cysteine 184.

## Discussion

Senescence is characterized by an irreversible arrest in cell proliferation(Hayflick, 1965). Cells undergo senescence because of a myriad of stresses, including DNA damage, mitochondrial damage, and oncogene overexpression(Bihani et al., 2007, 2004; Casar et al., 2018; Chen and Ames, 1994; Chen et al., 1998; Coppé et al., 2008; D’Adda Di Fagagna, 2008; D’Adda Di Fagagna et al., 2003; Di Leonardo et al., 1994; Franza et al., 1986; Land et al., 1983; Robles and Adami, 1998; Serrano et al., 1997; Wiley et al., 2016; Woods et al., 1997). Senescent cells exhibit a multi-faceted physiological response, where they exhibit a flattened morphology, increase in cell size(Chen and Ames, 1994; Serrano et al., 1997), upregulation of tumor suppressor proteins(Calabrese et al., 2009; Lowe et al., 2004; Stein et al., 1990; Zindy et al., 2003), expression of neutral pH active β-galactosidase (SA β-gal)(Dimri et al., 1995; Lee et al., 2006), and altered metabolic state(Bittles and Harper, 1984; Jones et al., 2005; Wiley and Campisi, 2021, 2016; Zwerschke et al., 2003). Arachidonic acid metabolism is upregulated in senescent cells, which leads to increased synthesis of eicosanoid prostaglandins, which regulate the physiology of senescent cells(Wiley et al., 2021; Wiley and Campisi, 2021). Senescent cells exhibit a secretory phenotype (SASP) consisting of a variety of bioactive molecules including cytokines and chemokines, growth factors, matrix metalloproteases, etc(Coppé et al., 2008). Senescent cells influence the surrounding cells via the SASP factors, which regulate proliferation, migration, and other cell biological processes in the neighboring cells(Campisi, 2005). SASP-mediated perturbations in the microenvironment are implicated in several senescence-associate pathologies(Wiley and Campisi, 2021). Senescent fibroblasts increase the proliferation of premalignant and malignant epithelial cells(Krtolica et al., 2001). Conditioned medium of senescent fibroblasts promoted tumorigenesis in mouse keratinocytes(Dilley et al., 2003). Senescent fibroblasts transform pre-malignant breast cancer cells into invasive, tumor-forming cells(Parrinello et al., 2005). Senescence in muscle stem cells induces sarcopenia via activation of the p38 MAP kinase pathway and transient inhibition of the p38 MAP kinases rejuvenates aged muscle stem cells to ameliorate sarcopenia(Cosgrove et al., 2014). Senescent cells inhibit the differentiation of myoblasts by secretion of IL6 by senescent muscle stem cells in myotonic dystrophy(Conte et al., 2023).

In this study, we show that senescent myoblasts synthesize and release eicosanoid prostaglandin 15-deoxy-Δ^12,14^-prostaglandin J_2_ (15d-PGJ_2_) (Fig. 1I and J), the terminal non-enzymatic dehydration product of prostaglandin PGD_2_(Shibata et al., 2002). We used Doxorubicin (Doxo) to induce senescence in C2C12 myoblasts and showed that the conditioned medium of senescent C2C12 cells inhibits differentiation of C2C12 myoblasts (Fig. 2A). Inhibition of synthesis of PGD_2_ by treatment of senescent cells with AT-56, a well-characterized inhibitor of prostaglandin D synthase(Hu et al., 2021; S. Hu et al., 2023; Shunfeng Hu et al., 2023; Irikura et al., 2009), rescued this inhibitory effect of the conditioned medium on the differentiation of myoblasts (Fig. 2A). A study has shown that prostaglandin PGD_2_ inhibits differentiation of C2C12 myoblasts(Veliça et al., 2010), but the authors noted that knockout of DP1 and DP2 (the known receptors of prostaglandins PGD_2_) does not abrogate inhibition of differentiation of myoblasts by PGD_2_. This observation suggested that PGD_2_ might inhibit the differentiation of myoblasts by a receptor-independent mechanism, possibly by its spontaneous non-enzymatic dehydration to 15d-PGJ_2_. 15d-PGJ_2_ has been suggested to be an endogenous ligand of PPARγ(Li et al., 2019). However, the inhibition of PPARγ did not abrogate the inhibition of differentiation of C2C12 myoblasts by 15d-PGJ_2_, suggesting the existence of other possible mechanisms(Hunter et al., 2001). 15d-PGJ_2_ has varied effects on cell physiology in a context-dependent manner. On one hand, 15d-PGJ_2_ promotes tumorigenesis by inducing epithelial to mesenchymal transition in breast cancer cell line MCF7(Choi et al., 2020), 15d-PGJ_2_ inhibits the proliferation of A549, H1299, and H23 lung adenocarcinoma cells via induction of ROS and activation of apoptosis(Slanovc et al., 2024). Here, we show that 15d-PGJ_2_ inhibits the proliferation and the differentiation of C2C12 myoblasts (Fig. 2B, C and D).

15d-PGJ_2_ contains an electrophilic cyclopentenone ring in its structure, allowing 15d-PGJ_2_ to covalently modify and form Michael adducts with cysteine residues of proteins(Shibata et al., 2002). A previous proteomic study in endothelial cells showed biotinylated 15d-PGJ_2_ covalently modified over 300 proteins, which regulate several physiological processes including cell cycle (MAPK1, MCM4), cell metabolism (Fatty acid synthase, Isocitrate dehydrogenase), apoptosis (PDCD6I), translation (Elongation factor 1 and 2, EIF4A-I), intracellular transport (Importin subunit β1, Exportin 2, Kinesin 1 heavy chain)(Marcone and Fitzgerald, 2013). Another proteomic study in neuronal cells suggested that 15d-PGJ_2_ modifies several proteins including chaperone HSP8A, glycolytic proteins Enolase 1 and 2, GAPDH, PKM1, cytoskeleton proteins Tubulin β2b, β actin, GFAP, etc(Yamamoto et al., 2011). This study also showed modification of peptide fragments homologous to IκB kinase β, Thioredoxin, and a small molecule GTPase HRas. 15d-PGJ_2_ has been shown to covalently modify HRas in NIH3T3 and Cos1 cells(Luis Oliva et al., 2003) and IMR90 cells(Wiley et al., 2021). Modification by 15d-PGJ_2_ led to the activation of HRas, judged by an increase in GTP-bound HRas. It is clear that 15d-PGJ_2_ is capable of modifying numerous proteins in different contexts. Despite these observations, the functional relevance of these modifications in numerous contexts remains to be mapped. Here we focused on the role of 15d-PGJ_2_ in the context of senescence and skeletal muscle differentiation. In this study, we showed that 15d-PGJ_2_ covalently modifies HRas at cysteine 184 but not cysteine 181 in C2C12 myoblasts (Fig. 3A). We showed by FRET microscopy that modification of HRas by 15d-PGJ_2_ in HRas WT and HRas C181S activates HRas in C2C12 cells, but 15d-PGJ_2_ is unable to activate HRas C184S in this context (Fig. 3B, C, and D and Fig. S3A and B). This observation shows a direct link between the modification of HRas by 15d-PGJ_2_ and the activation of HRas GTPase.

HRas activates two major downstream signaling pathways, the HRas-MAPK and the HRas-PI3K pathway(Pylayeva-Gupta et al., 2011). We showed that covalent modification of HRas by 15d-PGJ_2_ via the electrophilic cyclopentenone ring activates HRas (Fig. 3C and D) and activates the HRas-MAPK pathway, demonstrated by an increase in the phosphorylation of Erk after treatment with 15d-PGJ_2_ (Fig. 3E and F). However, we did not observe activation of the HRas-PI3K pathway, as we did not see an increase in the phosphorylation of Akt after treatment with 15d-PGJ_2_ (Fig. S3C). MAPK and PI3K pathways are known regulators of muscle differentiation(Bennett and Tonks, 1997; Rommel et al., 1999), where inhibition of the RAF-MEK-Erk pathway or activation of the PI3K pathway promotes the differentiation of myoblasts. Preferential activation of the HRas-MAPK pathway over the HRas-PI3K pathway after treatment with 15d-PGJ_2_ can be a possible mechanism by which 15d-PGJ_2_ can inhibit the differentiation of myoblasts. HRas is known to regulate the differentiation of myoblasts in different contexts. Constitutively active HRas signaling by expression of oncogenic HRas mutant (HRas V12) leads to inhibition of differentiation of myoblasts(Konieczny et al., 1989; Lassar et al., 1989; Olson,’ et al., 1987; Van Der Burgt et al., 2007). Here we showed that the inhibition of differentiation of myoblasts after 15d-PGJ_2_ is partially rescued in cells expressing the C184S mutant of HRas but not the wild type or the C181S mutant (Fig. 4D and E and S4E). HRas C184S did not get modified by 15d-PGJ_2_ (Fig. 3A). These observations suggest that the inhibition of differentiation of myoblasts by 15d-PGJ_2_ is partially dependent on the covalent modification of HRas by 15d-PGJ_2_.

Cysteine 181 and 184 in the C-terminal of HRas regulate the intracellular distribution of HRas between the plasma membrane and the Golgi by reversible palmitoylation and de-palmitoylation(Rocks et al., 2005). Inhibition of the palmitoylation of C-terminal cysteine 181, either by treatment with protein palmitoylation inhibitor 2-bromopalmitate or mutation of cysteine to serine, leads to accumulation of HRas at the Golgi. Intracellular localization of HRas maintains two distinct pools of HRas activity, where the plasma membrane pool shows a faster activation followed by short kinetics and the Golgi pool shows a slower activation but a sustained activation(Agudo-Ibáñez et al., 2015; Busquets-Hernández and Triola, 2021; Lorentzen et al., 2010; Rocks et al., 2005). We showed that the covalent modification of HRas by 15d-PGJ_2_ alters the intracellular distribution of HRas. We showed that the covalent modification of HRas by 15d-PGJ_2_ leads to an increase in the localization of the wild-type HRas at the plasma membrane compared to the Golgi (Fig. 4B). We did not observe any changes in the intracellular distribution of HRas C181S or HRas C184S after treatment with 15d-PGJ_2_ (Fig. 4C). HRas C184S is not modified by 15d-PGJ_2_, but HRas C181S is modified by 15d-PGJ_2_ (Fig. 3A). This suggests that the intracellular redistribution of HRas due to covalent modification by 15d-PGJ_2_ at cysteine 184 requires palmitoylation of cysteine 181.

Previous reports suggest that downstream signaling of HRas depends on the intracellular localization of HRas(Rocks et al., 2005; Santra et al., 2019). For example, targeted localization of HRas at the ER membrane induced expression of cell-migration genes. Localization of HRas at the plasma membrane showed a strong correlation with the expression of cell cycle genes, particularly the MAPK signaling pathway. Localization of HRas at the plasma membrane also showed a negative correlation with genes associated with the PI3K-Akt pathway. Here we showed that the intracellular distribution of HRas regulates differentiation of myoblasts. In order to show this, we used the constitutively active mutant of HRas (HRas V12) which has been shown to inhibit the differentiation of myoblasts(Engler et al., 2021; Konieczny et al., 1989; Lassar et al., 1989; Olson,’ et al., 1987; Scholz et al., 2009; Van Der Burgt et al., 2007). We expressed cysteine mutants of HRas V12 in C2C12 myoblasts and found that HRas V12 C181S localized predominantly at the Golgi whereas HRas V12 and HRas V12 C184S localized at both the plasma membrane and the Golgi (Fig. S4A). When differentiated, we observed that C2C12 cells expressing HRas V12 C181S differentiated but HRas V12 or HRas V12 C184S did not differentiate (Fig. S4B, C, and D). These observations suggest alteration of intracellular distribution of HRas affects the HRas-mediated inhibition of the differentiation of myoblasts.

Doxorubicin (Doxo) is a widely used chemotherapy agent for the treatment of cancers(Johnson-Arbor and Dubey, 2022). Treatment with Doxo induces senescence. Doxo-mediated DNA damage leads to p53, p16, and p21-dependent senescence in human fibroblasts(Di Leonardo et al., 1994; Robles and Adami, 1998). On the other hand, treatment with doxorubicin leads to a decrease in muscle mass and cross-sectional area, leading to chemotherapy-induced cachexia(Hiensch et al., 2020). Several mechanisms have been proposed behind chemotherapy-induced cachexia, including the generation of reactive oxygen species(Gilliam and St. Clair, 2011), activation of proteases like calpain and caspases(Gilliam et al., 2012; Smuder et al., 2011), and impaired insulin signaling(de Lima Junior et al., 2016). This study provides a possible mechanism behind chemotherapy-induced loss of muscle mass and functioning. Induction of senescence in myoblasts by treatment with Doxo could lead to increased synthesis and release of 15d-PGJ_2_ by senescent cells which could be taken up by myoblasts in the microenvironment. The lipid could covalently modify and activate HRas at cysteine 184 to inhibit the differentiation of myoblasts. Therefore, targeting the synthesis and release of 15d-PGJ_2_ by senescent cells could serve as an important target to promote skeletal muscle homeostasis in cancer patients.

## Supplementary Information

**Figure S1.**
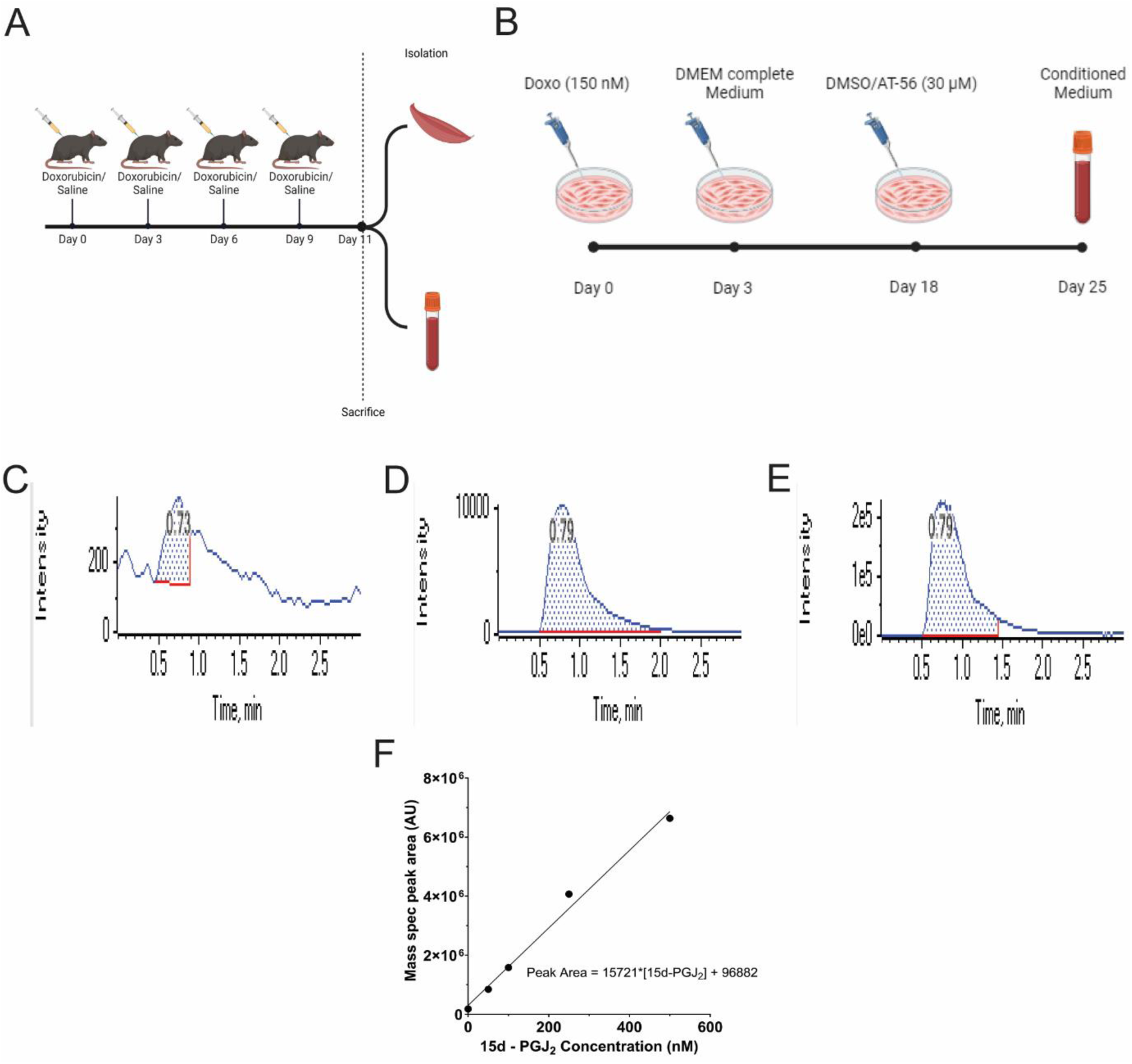
Treatment with Doxo induces senescence *in vivo* and *in vitro* and induces release of eicosanoid prostaglandin 15d-PGJ_2_. A. Schematic Representation of the experimental flow for treatment of B6J mice with Doxo (5 mg/kg) or Saline. B. Schematic representation of the experimental flow for treatment of senescent C2C12 cells with prostaglandin D synthase (PTGDS) inhibitor AT-56 (30 µM) or DMSO. C. Representative peak of quantification of m/z 315.100 → m/z 271 transitions from blank samples. D. Representative peak of quantification of m/z 315.100 → m/z 271 transitions from conditioned medium of C2C12 myoblasts treated with DMSO. E. Representative peak of quantification of m/z 315.100 → m/z 271 transitions from conditioned medium of C2C12 myoblasts treated with Doxo (150 nM). _F._ Standard curve of m/z 315.100 → m/z 271 fragment peak areas vs concentrations of 15d-PGJ_2_

**Figure S2.**
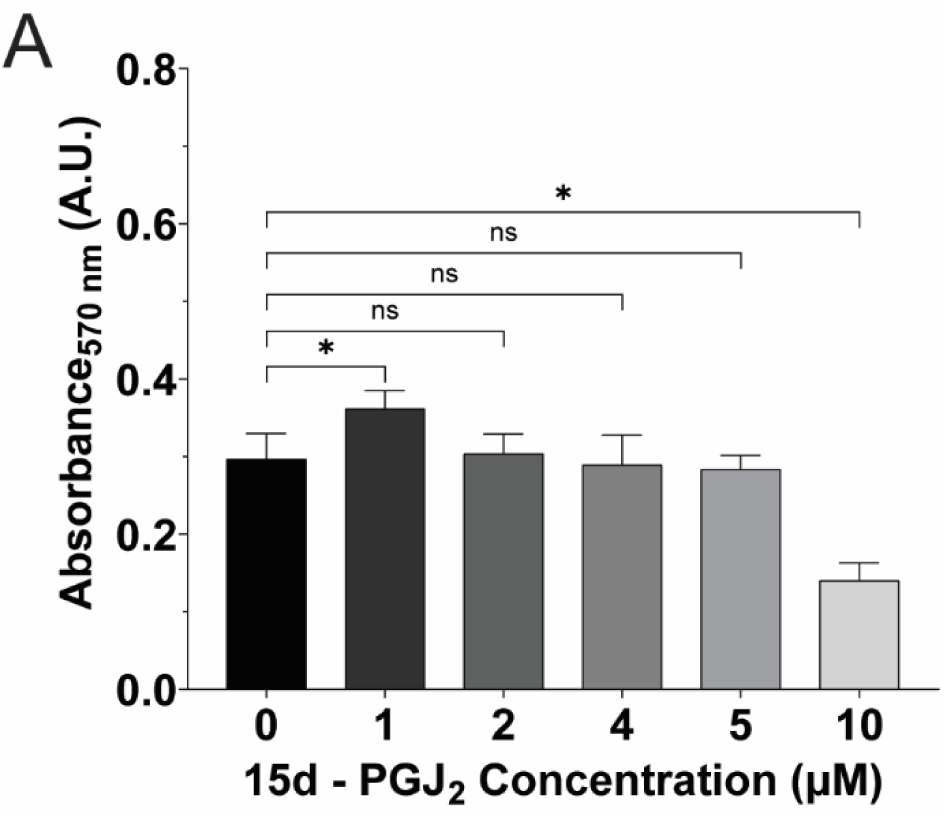
Viability of C2C12 myoblasts after treatment with 15d-PGJ_2_ (10 µM) in differentiating medium. A. Viability of C2C12 cells, measured by MTT assay, after 24 hours of treatment with 15-PGJ_2_ (0 µM, 1 µM, 2 µM, 4 µM, 5 µM, 10 µM). (Statistical significance was tested by the two-tailed student’s t-test ns=p>0.05, *=p<0.05, **=p<0.01, ***=p<0.001, ****=p<0.0001)

**Figure S3.**
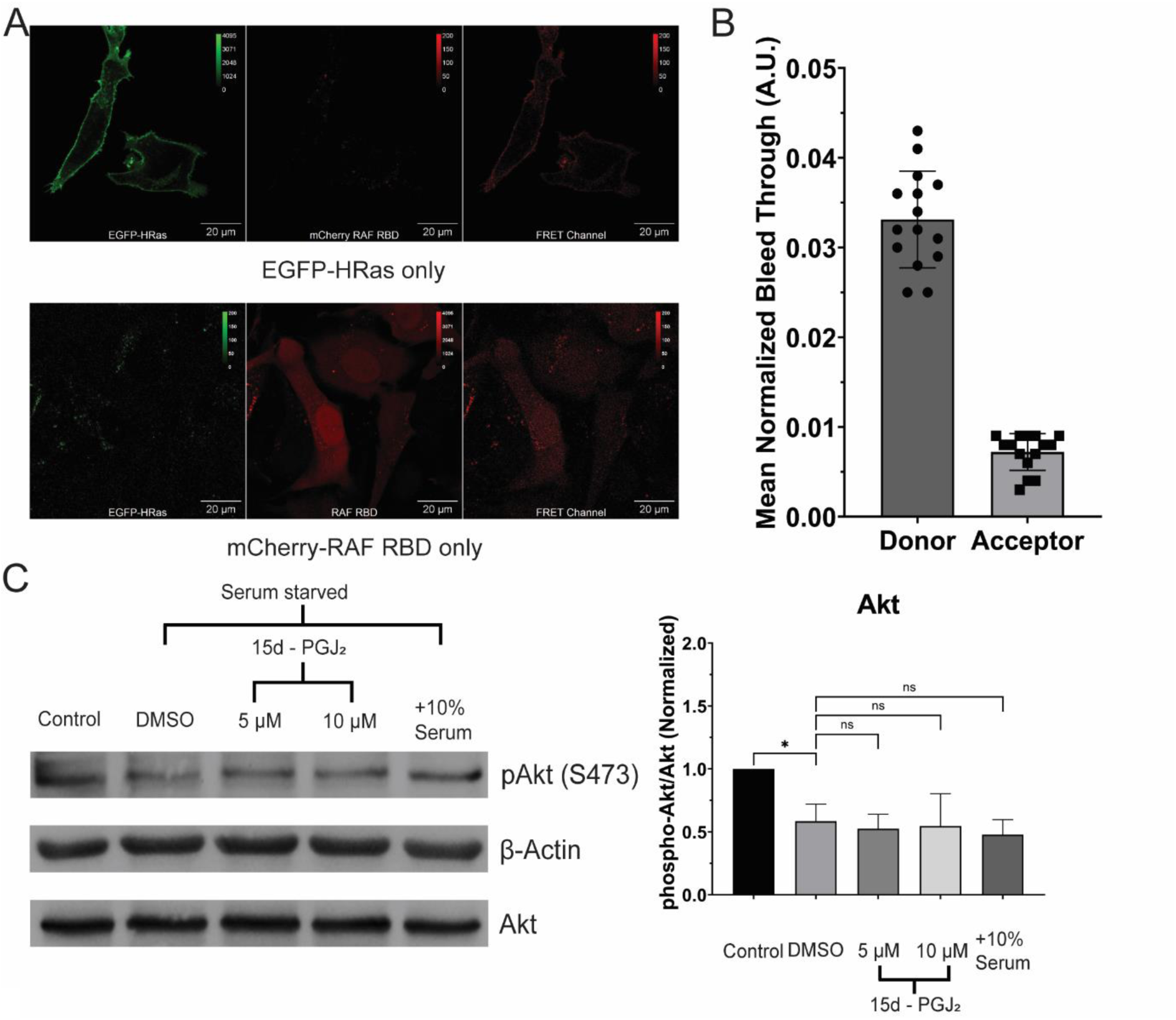
Activation of HRas after treatment with 15d-PGJ_2_. A. Representative confocal micrograph of C2C12 cells expressing only EGFP-tagged HRas or mCherry-tagged RAF RBD alone for spectral overlap (bleed-through) calculations. B. Mean bleed-through of EGFP-tagged HRas (Donor) and mCherry-tagged RAF RBD (Acceptor) in the FRET channel. C. Phosphorylation of Akt (measured by immunoblotting) in C2C12 cells treated with 15d-PGJ_2_ (5, 10 µM) or DMSO for 1 hr after starving the cells in 0.2% serum medium for 24 hrs. The densitometric ratio of phosphorylated Akt (Ser473) to total Akt was normalized to non-starved C2C12 cells. (Statistical significance tested by two-tailed heteroscedastic student’s t-test, N=3). (Statistical significance was tested by the two-tailed student’s t-test ns=p>0.05, *=p<0.05, **=p<0.01, ***=p<0.001, ****=p<0.0001)

**Figure S4.**
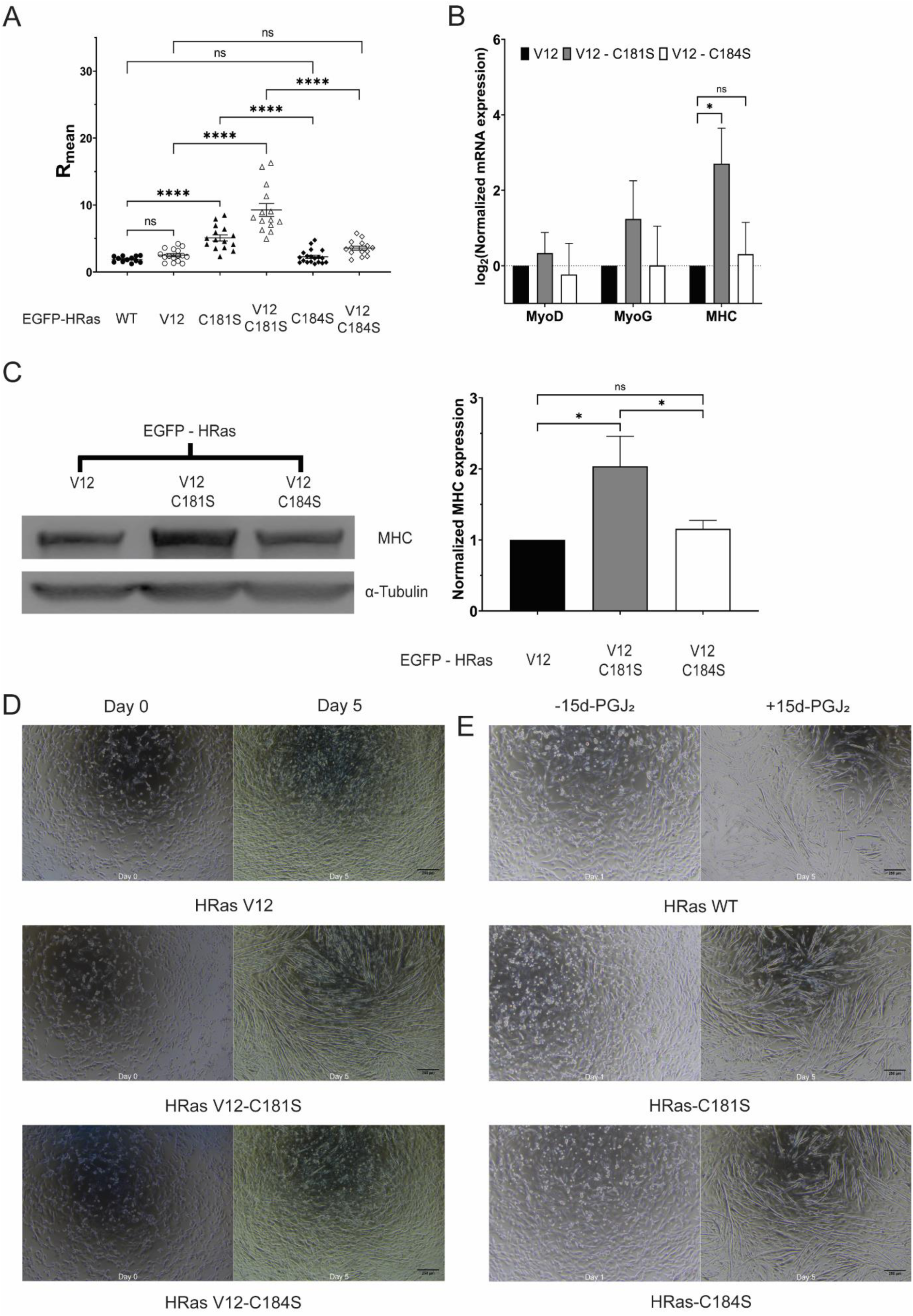
C-terminal cysteine-mediated intracellular distribution of constitutively active HRas (HRas V12) regulates the differentiation of C2C12 myoblasts. A. Distribution of EGFP-tagged HRas WT/ HRas V12/ HRas-C181S/ HRas V12-C181S/ HRas-C184S/ HRas V12-C184S between the Golgi complex and the plasma membrane, as seen by the scatter plot of R_mean_, the ratio of the mean HRas intensity at the Golgi complex to that at the plasma membrane, in C2C12 cells. (Statistical significance tested by two-tailed heteroscedastic student’s t-test, N=3) B. mRNA levels of MyoD, MyoG, and MyHC relative to 18s rRNA (measured by quantitative PCR) in C2C12 cells expressing EGFP-tagged HRas V12/ HRas V12-C181S/ HRas V12-C184S in C2C12 differentiation medium for 5 days. (Statistical significance tested by two-tailed heteroscedastic student’s t-test, N=3) C. Protein levels of MyHC (measured by immunoblotting) in C2C12 cells expressing EGFP-tagged HRas V12/HRas V12-C181S/HRas V12-C184S in C2C12 differentiation medium for 5 days. The densitometric ratio of levels of MyHC to α-Tubulin was normalized to C2C12 cells expressing HRas V12. (Statistical significance tested by two-tailed heteroscedastic student’s t-test, N=3). D. Brightfield image of C2C12 cells expressing EGFP-HRas V12, V12 C181S, or V12 C184S on Day 0 and Day 5 of differentiation. E. Brightfield image of C2C12 cells expressing EGFP-HRas WT, C181S, or C184S after 5 days of treatment with 15d-PGJ_2_ or DMSO. (Statistical significance was tested by the two-tailed student’s t-test ns=p>0.05, *=p<0.05, **=p<0.01, ***=p<0.001, ****=p<0.0001)

**Supplementary Table 1:** 15d-PGJ_2_ conc. (pmol and fg/cell) detected in samples by mass spectrometry.

| Sample | 15d-PGJ<sub>2</sub><br>(pmol) | Cell<br>Number | 15d-PGJ<sub>2</sub>/Cell<br>(fg) |
| --- | --- | --- | --- |
| Conditioned medium Quiescent C2C12 cells_1 | 27.087 | 462500 | 18.531 |
| Conditioned medium Quiescent C2C12 cells_2 | 18.080 | 650000 | 8.801 |
| Conditioned medium Senescent C2C12 cells_1 | 809.633 | 287500 | 891.018 |
| Conditioned medium Senescent C2C12 cells_2 | 940.032 | 300000 | 991.420 |

**Supplementary Table 2:**
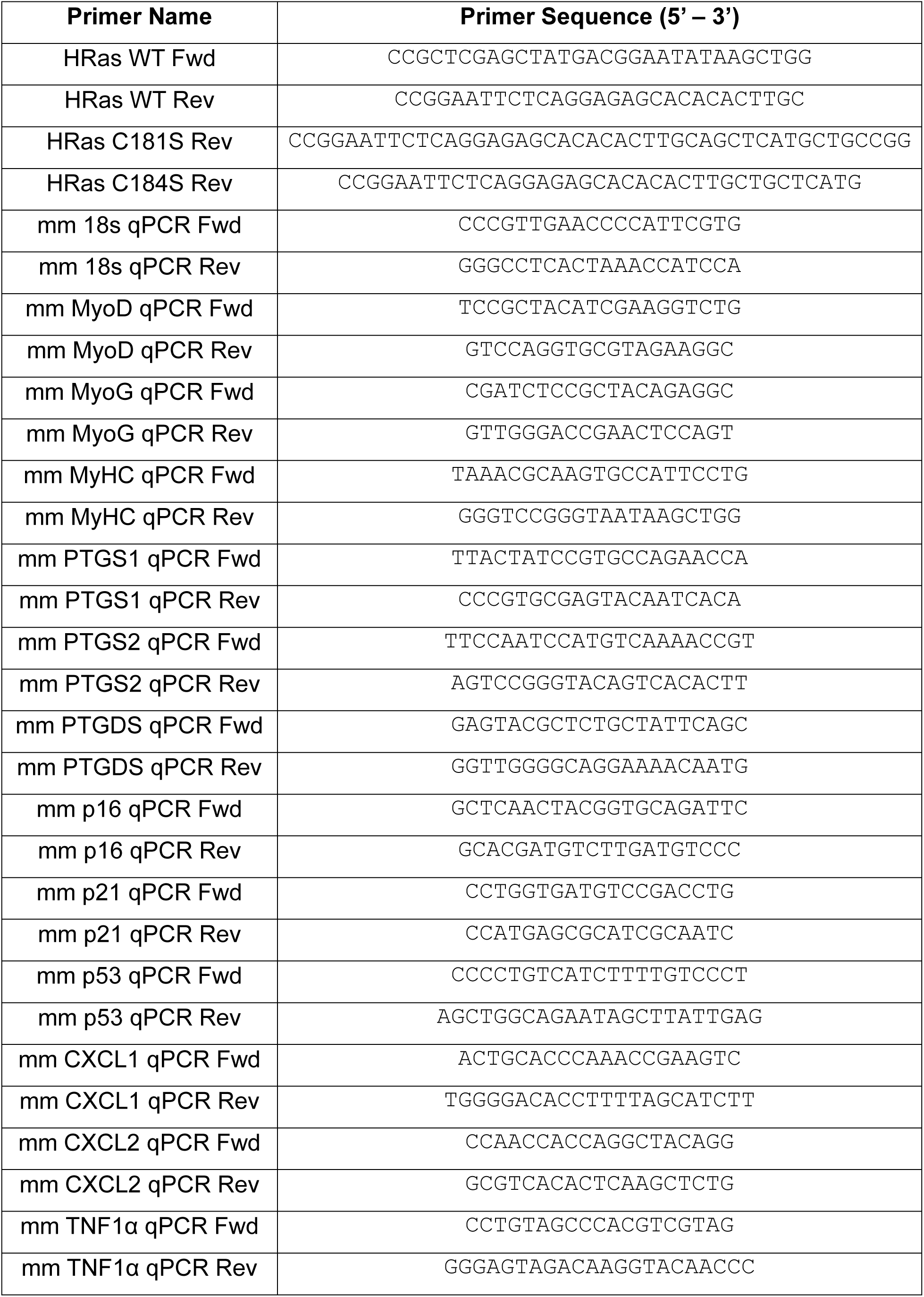
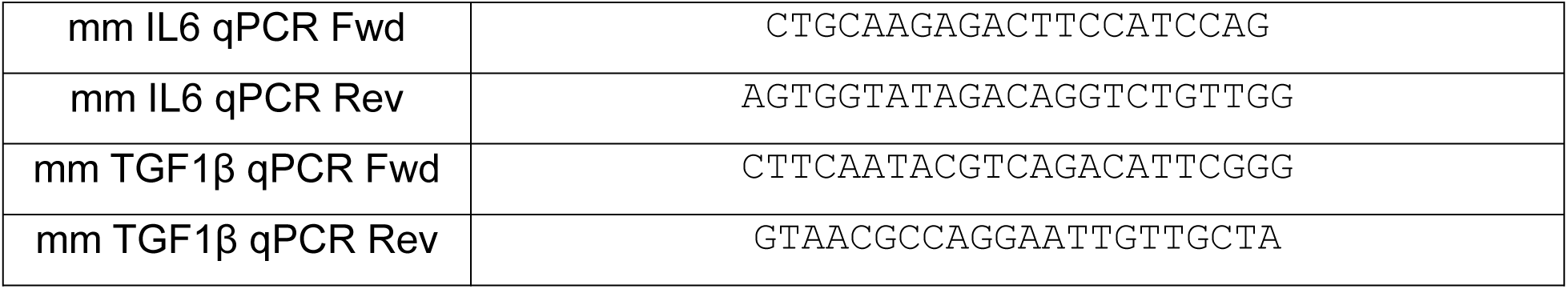
List of Primers:

**Supplementary Table 3:** List of Reagents.

| Item Description | Manufacturer | Catalog No. |
| --- | --- | --- |
| pEGFP-C1 | Clontech |  |
| Phusion high fidelity DNA polymerase | Thermo Scientific | F 530 |
| Xho1 restriction enzyme | New England Biolabs Inc. | R0146 |
| EcoR1 restriction enzyme | New England Biolabs Inc. | R0101 |
| T4 DNA Ligase | Takara Bio | 2011 |
| DMEM High Glucose | Gibco | 11995065 |
| FBS, Certified (Origin: United States) | Gibco | 16000044 |
| Horse Serum, Heat Inactivated (Origin: New Zealand) | Gibco | 26050070 |
| Penicillin – Streptomycin – Glutamine (100x) | Gibco | 10378016 |
| Penicillin – Streptomycin (100x) | Gibco | 15140163 |
| DPBS, no calcium, no magnesium | Gibco | 14190144 |
| 0.25% Trypsin - EDTA | Gibco | 25200056 |
| jetPRIME transfection reagent | Polyplus | 101000046 |
| Doxorubicin | Sigma – Aldrich | D1515 |
| 15-deoxy-$\Delta^{12,14}$-Prostaglandin J<sub>2</sub> | Cayman Chemical Company | 18570 |
| 9,10-dihydro-15-deoxy-$\Delta^{12,14}$-Prostaglandin J<sub>2</sub> | Cayman Chemical Company | 18590 |
| 15-deoxy-$\Delta^{12,14}$-Prostaglandin J<sub>2</sub>-Biotin | Cayman Chemical Company | 10141 |
| Dynabeads™ MyOne™ Streptavidin C1 | Invitrogen | 65001 |
| cOmplete Protease inhibitor cocktail tablets | Roche | 11697498001 |
| Wheat germ agglutinin, Alexa fluor 633 conjugate | Invitrogen | 2307289 |
| ProLong Gold Antifade Mounting medium | Invitrogen | P36930 |
| Paraformaldehyde | Sigma – Aldrich | 158127 |
| TRIzol reagent | Invitrogen | 15596018 |
| PrimeScript™ 1st strand cDNA Synthesis Kit | Takara Bio | 6110A |
| PowerUp™ SYBR™ Green Master Mix | Applied Biosystems | A25742 |
| WesternBright™ ECL-spray Western blotting detection system | advansta | K-12049-D50 |
| Fetuin (Bovine) | Sigma – Aldrich | F2379 |
| hEGF | Sigma – Aldrich | E9644 |
| N2 Supplement | Thermo Scientific | 17502048 |
| Dexamethasone | Sigma – Aldrich | D4902 |
| DMEM Low Glucose | Thermo Scientific | 10567014 |
| Bovine Serum Albumin Fraction-V, Cell culture tested | HIMEDIA | 9048468 |
| Pierce™ Streptavidin Magnetic Beads | Thermo Scientific | 88816 |

**Supplementary table 4:** List of Antibodies.

| **Antibody Name** | **Clonality** | **Species of Origin** | **Manufacturer** | **Catalog No.** |
| --- | --- | --- | --- | --- |
| Phospho-Erk (Thr202/<br>Tyr204) | Polyclonal | Rabbit | Cell Signaling<br>Technology | 9101S |
| Erk | Polyclonal | Rabbit | Cell Signaling<br>Technology | 9102S |
| GAPDH | Monoclonal | Mouse | Puregene | PG23002 |
| β-actin | Polyclonal | Rabbit | Cell Signaling<br>Technology | 4967S |
| Phospho-Akt (Ser473) | Monoclonal | Mouse | Cell Signaling<br>Technology | 4051S |
| Akt | Monoclonal | Rabbit | Cell Signaling<br>Technology | 4691S |
| GFP | Monoclonal | Rabbit | Cell Signaling<br>Technology | 2956S |
| Myosin Heavy Chain | Monoclonal | Mouse | Invitrogen | 14-6503-82 |
| Myosin Heavy Chain | Monoclonal | Mouse | Developmental<br>Studies Hybridoma<br>Bank (DSHB) | MF20 |
| p21 | Monoclonal | Mouse | Santacruz<br>Biotechnology | sc-6246 |
| γH2A.X | Polyclonal | Rabbit | Novus Biologicals | NB100384 |
| Tubulin | Monoclonal | Mouse | Cell Signaling<br>Technology | 3873S |
| HRP - Anti-Mouse |  | Horse | Cell Signaling<br>Technology | 7076S |
| HRP - Anti-Rabbit |  | Goat | Cell Signaling<br>Technology | 7074P2 |
| Alexa Fluor 568 – Anti-Mouse | Polyclonal | Goat | Invitrogen | A11031 |

## Materials and Methods

### Plasmids

Unmutated and cysteine mutants of HRas WT [HRas WT, HRas-C181S, and HRas-C184S] and HRas V12 [HRas V12, HRas V12-C181S, HRas V12-C184S] were cloned in the pEGFPC1 vector (Clontech) by restriction digestion-ligation method. Constructs of wild-type HRas were PCR amplified from a previously available HRas construct in the lab with construct-specific primers. Proper nucleotide additions were made to the forward primer to maintain the EGFP ORF, marking a 7 amino acid linker between the proteins. The construct sequences were confirmed by Sanger sequencing. GalT-TagRFP construct was a gift from Prof. Satyajit Mayor and was used to mark the Golgi. mCherry-RAF-RBD construct was a gift from Prof. Phillipe Bastiens and was used to measure the activity of HRas GTPase using FRET.

### Cell Maintenance

C2C12 mouse myoblasts (CRL–1772) were obtained from ATCC and were maintained in DMEM complete medium @ 37° C, 5% CO_2_. For experiments, the cells were trypsinized with 0.125% trypsin-EDTA (Gibco) and were seeded in required numbers in cell culture dishes. Human Skeletal Muscle Myoblast (CC-2580) were obtained from Lonza and were maintained in DMEM Skeletal Muscle growth medium @ 37° C, 5% CO_2_. For experiments, the cells were trypsinized with 0.125% trypsin – EDTA (Gibco) and were seeded in required numbers in cell culture dishes. All cultures tested negative for mycoplasma checked by Mycoalert Mycoplasma Detection Kit (Lonza).

### Conditioned media collection

C2C12 cells seeded in 60mm dishes were treated with Doxorubicin (150 nM) for 3 days. The media was then changed to DMEM complete medium without Doxorubicin for 19 days after treatment with Doxorubicin. The cells were treated with DMSO or AT-56 (30 µM) in the DMEM complete medium for 2 days. On Day 21, the cells were treated with DMSO or At-56 in DMEM Starvation medium for 3 days. The media was then collected and centrifuged @1000g, R.T. for 5 minutes. The media was then stored at -80° C after flash freezing in liq. N_2_ till further requirement.

### Treatments

15d-PGJ_2_ (Cayman Chemical Company) dissolved in DMSO (10 mM) was diluted in DMEM media for experiments. 9,10-dihydro-15d-PGJ_2_ (Cayman Chemical Company) dissolved in DMSO (10 mM) was appropriately diluted in DMEM media for experiments. DMSO was used as vehicle control. A media change of the same composition was given every 24 hours. C2C12 cells with 70-80% confluency were treated with Doxorubicin (Doxo) for 3 days. After 72 hours, Doxo was removed from the medium and the cells were kept for 10 more days with media change every 3 days till the end of the experiment. C2C12 cells transfected with EGFP-HRas WT/C181S/C184S in 35mm dishes were treated with 15d-PGJ_2_-Biotin (5 µM) in DMEM Hi Glucose medium (Gibco) supplemented with 1% Penicillin-streptomycin-Glutamine (Gibco) without fetal bovine serum for 3 hours. Conditioned medium collected from senescent cells was thawed @ 37° C. The medium was then supplemented with 2% heat-inactivated horse serum and 1% penicillin-streptavidin-glutamine. C2C12 myoblasts were treated with the conditioned medium and were given a media change every 48 hours.

### Transfections

C2C12 cells were seeded in 35 mm dishes to achieve confluency of ∼60-70%. For western blot, immunoprecipitation, and differentiation experiments, the cells were transfected with EGFP-tagged HRas WT/ HRas-C181S/ HRas-C184S/ HRas V12/ HRas V12-C181S/ HRas V12-C184S using the jetPRIME transfection reagent (Polyplus) using the manufacturer’s protocol. For measuring the intracellular distribution of HRas, the cells were reverse transfected with EGFP-tagged HRas WT/ HRas-C181S/ HRas-C184S/ HRas V12/ HRas V12-C181S/ HRas V12-C184S and GalT-TagRFP, a Golgi apparatus marker protein tagged with red fluorescent TagRFP protein using the jetPRIME transfection reagent using the manufacturer’s protocol. For measuring the activity of HRas, the cells seeded in imaging dishes (iBidi) were transfected with EGFP-HRas and mCherry-RAF-RBD using jetPRIME transfection reagent (Polyplus) using the manufacturer’s protocol. Transfection efficiency was confirmed by checking for GFP and RFP fluorescence after 24 hours of transfection.

### Myoblast differentiation

C2C12 cells were treated with either 15d-PGJ_2_ or DMSO in the C2C12 differentiation medium. The cells were given a media change of the same composition every 24 hours. The cells were harvested after 5 days of 15d-PGJ_2_ treatment for either RNA or protein isolation. Human Skeletal Muscle Myoblast cells were treated with DMSO or 15d-PGJ_2_ in the Skeletal Muscle Differentiation medium. A media change of the same composition was given every 24 hours. The cells were harvested after 5 days of treatment for protein isolation.

### X-Gal staining

Proliferative and Doxo-treated C2C12 cells were fixed with 0.25% glutaraldehyde, washed with PBS, and incubated overnight in X-gal staining solution at 37° C in a CO_2_-free chamber. The presence of the Indigo blue product was confirmed using the Ti2 widefield inverted microscope (Nikon).

### Immunoprecipitation

C2C12 cells transfected with EGFP-HRas and treated with 15d-PGJ_2_-Biotin were harvested and lysed in RIPA-PP buffer and the lysate was centrifuged @15000 rpm, 4° C, 30 minutes. Protein estimation was done using the BCA assay kit (G Biosciences). 100 µg of protein was loaded on 10 µl MyOne Streptavidin C1 dynabeads blocked with 1% BSA in IP washing buffer. The lysate-streptavidin mix was incubated @4° C, 10 rpm overnight. The beads were then washed with IP washing buffer and then boiled in 20 µl Laemmlli buffer. 15 µl of the beads were loaded on 12% SDS-Polyacrylamide gel for detection of EGFP-HRas by immunoblotting using EGFP antibody.

### Western blotting

For measuring Erk/Akt phosphorylation in C2C12 cells were seeded in 35 mm dishes. 1x 35 mm dish was harvested in RIPA – PP the next day, while the rest were incubated in DMEM starvation medium @37° C. The cells were treated with 15d - PGJ_2_ after 24 hrs of starvation @37° C. The cells were harvested at 1 hour after treatment in RIPA-PP. Protein quantification was done using BCA assay (G Biosciences) using the manufacturer’s protocol. An equal mass of proteins was loaded onto a 12% SDS Polyacrylamide gel in Laemmlli buffer. The proteins were transferred onto a PVDF membrane and were probed with phospho-Erk/Erk antibodies for measuring Erk phosphorylation and with phospho-Akt/Akt antibodies for measuring Akt phosphorylation. For measuring the expression of Myosin heavy chain, C2C12 cells expressing EGFP-tagged HRas WT/ HRas-C181S/ HRas-C184S/ HRas V12/ HRas V12-C181S/ HRas V12-C184S or Human Skeletal Muscle Myoblasts were seeded in 35 mm dishes and were harvested in RIPA-PP after 5 days of differentiation. Protein quantification was done using BCA assay (G Biosciences) using the manufacturer’s protocol. An equal mass of proteins was loaded onto an 8% SDS Polyacrylamide gel in Laemmlli buffer. The proteins were transferred onto a PVDF membrane and were probed with Myosin Heavy Chain Antibody.

### qPCR

C2C12 cells, untransfected or expressing EGFP-tagged HRas WT/ HRas-C181S/ HRas-C184S and treated with DMSO/15d - PGJ_2_, or expressing EGFP-tagged HRas V12/ HRas V12-C181S/ HRas V12-C184S were lysed in TRIZol at the end of the experiment (Invitrogen). RNA was isolated from the lysate by the chloroform-isopropanol method using the manufacturer’s protocol. The RNA was quantified and 1.5 μg of RNA was used to prepare cDNA using PrimeScript 1st strand cDNA Synthesis Kit (Takara Bio) and random hexamer primer. Gene expression for differentiation markers was measured by qPCR using PowerUp™ SYBR™ Green Master Mix (Applied Biosystems) and previously reported qPCR primers (Wang et al., 2012). Relative gene expression was quantified using the ΔΔC_T_ method (Livak and Schmittgen 2001) with 18s rRNA as an internal loading control and DMSO vehicle as an experimental control.

### Immunofluorescence

C2C12 cells were seeded in 35 mm dishes (Corning) on glass coverslips (Blue Star) coated with 0.2% Gelatin (Porcine, Sigma Aldrich) and were fixed with the fixative solution at the end of the experiment. The cells were then permeabilized and blocked with the blocking solution and were then incubated with Myosin Heavy Chain antibody in the blocking solution overnight. The cells were then washed with 1x PBS, incubated with fluorophore tagged secondary antibody, and were mounted in Prolong gold antifade medium with DAPI (Invitrogen). The cells were then imaged under the FV3000 inverted confocal laser scanning microscope (Olympus-Evident) using appropriate lasers and detectors.

### Confocal microscopy for measuring HRas distribution between the Golgi and the plasma membrane

C2C12 cells expressing EGFP-tagged HRas WT/ HRas-C181S/ HRas-C184S + GalT-TagRFP were starved overnight in DMEM starvation medium and treated with DMSO or 15d-PGJ_2_ (10 µM) in DMEM complete medium for 24 hrs, with a medium change @ 12 hrs post-treatment. The cells were then fixed with the fixative solution @ R. T., washed with PBS, and stained for plasma membrane with Alexa Fluor 633 conjugated Wheat Germ Agglutinin (WGA-633) (Invitrogen). The cells were washed with PBS and were then mounted on glass slides in ProLong Gold Antifade Mounting medium (Invitrogen). C2C12 cells expressing EGFP-tagged HRas V12/ HRas V12-C181S/ HRas V12-C184S were also fixed with the fixative solution @R. T., washed with PBS, stained with WGA-633, and mounted on slides in Prolong gold antifade medium (Invitrogen). The cells were imaged with the FV3000 inverted confocal laser scanning microscope (Olympus-Evident) using appropriate lasers and detectors. Preliminary image processing was done using ImageJ (NIH), while batch analysis of HRas at the plasma membrane and the Golgi complex was done using a custom MATLAB script, where EGFP-HRas image was overlayed onto the GalT-TagRFP and WGA-633 image to obtain HRas localization at the Golgi complex and the Plasma Membrane respectively. A ratio of mean HRas intensity at the Golgi complex to that of at the Plasma membrane (R_mean_) was calculated and was used to compare HRas distribution between treatments.

### FRET confocal microscopy to measure the intracellular activity of HRas

C2C12 cells expressing EGFP-tagged HRas WT/HRas C181S/HRas C184S and mCherry-RAF-RBD were starved overnight in the DMEM starvation medium. The cells were imaged with the FV3000 inverted confocal laser scanning microscope (Olympus-Evident) using the following lasers and detectors:

1. Donor Channel: 488nm excitation, 510 (+/-) 20nm detection.
2. Acceptor Channel: 561nm excitation, 630 (+/-) 50nm detection.
3. FRET Channel: 488nm excitation, 630 (+/-) 50nm detection.

The cells were then treated with 15d-PGJ_2_ (10 µM) for 1 hour and were imaged using the same imaging parameters. C2C12 cells expressing EGFP-HRas or mCherry-RAF RBD only were used to calculate the bleed-through corrections (EGFP emission @ 630 (+/-) 50nm, and Excitation of mCherry by 488 nm laser). Preliminary processing was done using ImageJ (NIH). The FRET index was calculated using the FRET and co-localization analyzer plugin(Hachet-Haas et al., 2006). The FRET index was then divided by the intensity of the Acceptor channel to normalize the variation in the expression of mCherry. We used the mean normalized FRET index to compare the activity of HRas before and after treatment with 15d-PGJ_2_.

### Quantification of myotube fusion index

Differentiated C2C12 myoblasts were immunostained for MHC and DAPI and were imaged on the FV3000 inverted confocal laser scanning microscope (Olympus-Evident). Analysis of the fusion index was done using the Myotube Analyzer Software(Noë et al., 2022). DAPI and MHC images were thresholded to remove background noise. The images were converted to binary masks and the channels were overlayed to obtain the no. of nuclei overlaying with MHC^+ve^ fibers. The fusion index was calculated as the percentage ratio of no. nuclei overlaying the MHC^+ve^ fibers to the total no. of nuclei in the field of view.

### Quantification of cell doubling time

Cells were counted every 24 hours and the normalization was done to the number of cells counted on day 0 of the treatment (to consider attaching efficiency and other cell culture parameters). Doubling time was calculated as the reciprocal of the slope of the graph of log2(normalized cell number) vs time.

### MTT Assay

An equal number of C2C12 cells were seeded in 96 well plates in replicates. MTT assay was done at the end of the experiment using the manufacturer’s protocol. MTT reagent (Sigma Aldrich) was dissolved in 1x DPBS (5 mg/ml) and was filter sterilized. MTT reagent was added to each well and the cells were incubated @37° C, 5% CO_2_ for 3 hours. The medium was removed at the end of the incubation and the precipitated crystals were dissolved in DMSO @37° C, 5% CO_2_ for 15 minutes. Absorbance @570 nm was recorded using the varioskan multimode plate reader (Thermo Scientific).

### Animal Experiments

Mice were maintained at BLiSC Animal Care and Resource Centre (ACRC). All the procedures performed were approved by the Internal Animal Users Committee (IAUC) and the Institutional Animal Ethics Committee (IAEC). 12–15-week-old C57BL/6J (JAX#000664) mice were injected intraperitoneally (I.P.) with 5 mg/kg Doxorubicin (Doxo) four times, once every three days. Intraperitoneal injection of Saline was used as a control. The mice were sacrificed on Day 11 after the first injection. Hindlimb muscles from 4 animals (control and treated with Doxo each) were used for qPCR analysis and Hindlimb muscles from 3 animals (control and treated with Doxo each) were used for immunohistochemical analysis.

### Lipid extraction and detection of 15d-PGJ_2_ by mass spectrometry

For lipid extraction, cell pellets were resuspended in 3ml of a methanol solvent [water: methanol: 2:1, 1% formic acid (FA)] whereas only 1 ml of methanol with 3% FA was added to the 2 ml of CM, making a uniform sample volume of 3 ml. Subsequently, 1 ml of ethyl acetate was added to each sample and mixed vigorously. Phase separation was done by centrifuging the mixture (12000xg, 4℃ for 10 mins), and the organic phase containing the lipid was collected. This process was repeated thrice in total and all the organic phases were combined and dried under a nitrogen stream at RT. The residues were resuspended in 100 µl of 50% acetonitrile in water with 0.1% FA and were subjected to mass spec analysis using the Waters® Acquity UPLC class I system The detection of 15d-PGJ_2_ was performed using an electrospray ionization source (ESI) operating in the negative ion mode and a quadrupole trap mass spectrometer (AB SCIEX QTRAP 6500) connected to a Waters® Acquity UPLC class I system (Waters, Germany) outfitted with a binary solvent delivery system with an online degasser and a column manager with a column oven coupled to a UPLC autosampler. 5 µl samples were injected into the union for analysis. Solvent A consisted of 0.1% ammonium acetate in water and solvent B was 0.1% ammonium acetate in a mixture of acetonitrile/water (95:5). For each run, the LC gradient was: 0 min, 20% B; 0.5 min, 20% B; 1.5 min, 90% B; 2.5 min, 20% B; 3min, 20% B. Analyte detection was performed using multiple reaction monitoring (MRM), 315.100 → 271.100 and 315.100 → 203.100. Source parameters were set as follows: capillary voltage 3.8 kV, desolvation gas flow 25 L/h, source temperature 350 °C, ion source gas 1 flow 40 L/h, and ion source gas 2 flow 40 L/h. Acquisition and quantification were completed with Analyst 1.6.3 and Multiquant 3.0.3, respectively (method adopted from (Morgenstern et al., 2018)). For the standards, 2ml media of different known concentrations (50nM, 100nM, 250nM, and 500nM) of 15d-PGJ_2_ were prepared and subjected to the same extraction procedure as that of CM. A standard curve was plotted with the known concentration and the mass spec peak area, and the concentration of the lipid in samples was calculated.

### Reagents

- DMEM complete medium: DMEM Hi Glucose medium (Gibco) supplemented with 1% Penicillin – Streptomycin – Glutamine (Gibco) and heat-inactivated 10% Fetal Bovine Serum (US origin) (Gibco).
- Basal Conditioned medium: DMEM Hi Glucose medium (Gibco) supplemented with 1% Penicillin – Streptomycin – Glutamine (Gibco) and heat-inactivated 2% Fetal Bovine Serum (US origin) (Gibco).
- C2C12 differentiation medium: DMEM Hi Glucose medium (Gibco) supplemented with 2% Horse Serum (Gibco) and 1% Penicillin – Streptomycin – Glutamine (Gibco).
- DMEM Starvation medium: DMEM Hi Glucose medium (Gibco) supplemented with 0.2%heat-inactivated fetal bovine serum (US origin) (Gibco) and 1% Penicillin – Streptomycin – Glutamine (Gibco).
- RIPA – PP buffer: RIPA buffer (Invitrogen) supplemented with protease inhibitor cocktail (Roche) and 5 mM Sodium Fluoride and 5 mM Sodium Orthovanadate.
- TBS – T buffer: 50 mM Tris-Cl (pH = 7.5), 150 mM NaCl and 0.1% Tween – 20 in water.
- PBS: 2.67 mM KCl, 1.47 mM KH_2_PO_4_, 137.93 mM NaCl, 8.06 mM Na_2_HPO_4_ in water.
- IP Washing Buffer: 150 mM NaCl, 0.1% SDS, 1% NP-40 in 50 mM Tris-Cl (pH=7)
- Fixative Solution: 4% (w/v) Paraformaldehyde (Sigma – Aldrich) in PBS.
- Blocking Solution: 2% Heat Inactivated FBS, 0.2% BSA, 0.2% Triton - X, 0.05% NaN_3_ in PBS.
- Skeletal Muscle Growth Medium: DMEM Low Glucose Medium (Gibco), supplemented with 1% Penicillin – Streptomycin – Glutamine (Gibco), heat-inactivated 10% Fetal Bovine Serum (US origin) (Gibco), Bovine Fetuin (50 µg/ml) (Sigma – Aldrich), Dexamethasone (0.4 µg/ml), and hEGF (10 ng/ml).
- Skeletal Muscle Differentiation Medium: DMEM low glucose medium (Gibco) supplemented with 2% Horse Serum, 1% Penicillin – Streptomycin (Gibco), and 1% N2 Supplement.

## Acknowledgments

We thank Prof. Satyajit Mayor (NCBS), Prof. Phillipe Bastiens, and Prof. Apurva Sarin (InStem) for providing the wid-type HRas construct, the mCherry-RAF RBD construct and the vector backbones respectively. We thank Dr. Neetu Saini (InStem) for her help with setting up the cell culture facility. We thank Mr. Heera Lal for his help with the animal work. We thank Dr. Kamlesh Kumar Yadav and Ms. Sudeshna Saha for their help during the project. We thank the Central Imaging and Flow Cytometry Facility (CIFF) (NCBS-InStem) for their support with microscopy. We thank the Animal Care and Resource Centre (ACRC) (NCBS-InStem) for their support with mouse experiments. We thank the Mass Spectrometry facility (NCBS-InStem) for their support with the mass spectrometry work.

## Author Contribution

SSP: Project conceptualization, Cell culture treatments and assays, Biochemistry, Microscopy, Image Processing, and analysis, Writing: original draft, review, and edits.

AB: Animal work, Cell culture treatments and assays, Biochemistry, Mass Spectrometry, Writing: review and edits.

AV: Cell culture treatments and assays, Image processing and analysis, writing: review and edits.

SSS: Cell culture treatments and assays, Biochemistry, Writing: review and edits. MAJ: Image processing and analysis.

RGHM: Mass Spectrometry.

AR: Project Conceptualization, Writing: original draft, review, and edits. Corresponding Author.

